# Multi-omics analysis reveals a crucial role for Retinoic Acid in promoting epigenetic and transcriptional competence of an *in vitro* model of human Pharyngeal Endoderm

**DOI:** 10.1101/2022.06.26.497457

**Authors:** Andrea Cipriano, Alessio Colantoni, Danielle Gomes, Mahdi Moqri, Alexander Parker, Matthew Caldwell, Francesca Briganti, Jonathan Fiorentino, Maria Grazia Roncarolo, Antonio Baldini, Katja G Weinacht, Gian Gaetano Tartaglia, Vittorio Sebastiano

## Abstract

*In vitro* differentiation of human Pluripotent Stem Cells (hPSCs) into different cell types has enabled the study of developmental processes that are impossible to dissect *in vivo*. This innovation has allowed for the derivation of therapeutically relevant cell types that can be used for downstream applications and studies. The Pharyngeal Endoderm (PE) is considered an extremely relevant developmental tissue since it acts as a precursor to a plethora of organ systems such as Esophagus, Parathyroids, Thyroids, Lung, and Thymus. While several studies have highlighted the importance of these cells, an *in vitro* platform to generate human PE cells is still missing. Here we fill this knowledge gap, by providing a novel *in vitro* protocol for the derivation of *bona fide* PE cells from hPSCs. We demonstrated that our PE cells robustly express Pharyngeal Endoderm markers, they are transcriptionally similar to PE cells isolated from *in vivo* mouse development and represent a transcriptionally homogeneous population. Importantly, we elucidated the contribution of Retinoic Acid in promoting a transcriptional and epigenetic rewiring of PE cells. In addition, we defined the epigenetic landscape of PE cells by combining ATAC-Seq and ChIP-Seq of histone modifications. The integration of these data led to the identification of new putative regulatory regions and to the generation of a gene regulatory network orchestrating the development of PE cells. By combining hPSCs differentiation with computational genomics, our work reveals the epigenetic dynamics that occur during human PE differentiation, providing a solid resource and foundation for research focused on the development of PE derivatives and modeling of their developmental defects in genetic syndromes.

## INTRODUCTION

Human embryogenesis is characterized by the progression of highly dynamic and temporal stages involving sequential chromatin and transcriptional changes, driven by extracellular and intracellular signaling pathways that occur in a cell type and stage-dependent manner (Resto Irizarry et al., 2020; Sonnen & Janda, 2021; Zheng et al., 2017). The proper regulation of these processes is essential for the accurate, robust, and reproducible development of progenitor-like cells into distinct cell types, forming a cooperative and cohesive network of physiological systems (Murry & Keller, 2008). Studying these processes is crucial since they provide insight that can be leveraged to identify key signaling pathways that coordinate human development, and to understand how their disruption contributes to developmental diseases (Magaletta et al., 2020; McDonald-McGinn et al., 2015; Motahari et al., 2019; Sonnen & Janda, 2021). Human Pluripotent Stem Cells (hPSCs) have greatly improved the ability to study human development and developmental related diseases, thanks to their capability to self-renew and differentiate into all cell types of the human body (Murry & Keller, 2008; Takahashi et al., 2007; Thomson et al., 1998) and the ease of derivation from genetically mutated somatic cells. However, in order to harness the full potentiality of this platform, it is essential to be able to mimic the signals that occur *in vivo* during hPSCs differentiation to direct the development of these cells to specific lineages.

One such lineage is the Pharyngeal Endoderm (PE), which contributes to the Pharyngeal apparatus (PA) in vertebrates. This structure is highly conserved among vertebrates and it is formed between E8.5-10.5 in mouse and E21-28 in human, with the contribution of cells from all three germ layers (Frisdal & Trainor, 2014; Graham & Richardson, 2012; Shone & Graham, 2014). The PE, which originates from the anterior-most region of the foregut, is considered the main driver orchestrating the development of the PA. This is primarily due to the formation of the Pharyngeal Pouches (PPs), valley-like structures within the PA, which emerge thanks to the out pocketing of the PE (Crump et al., 2004; Graham et al., 2005; Graham & Richardson, 2012; Veitch et al., 1999). The PPs serve as a microenvironment for physiological development and are essential for the morpho-patterning of important organs and structures such as the lining of the pharynx, palatine tonsils, inner ear, parathyroids, thyroid glands, ultimobranchial bodies, and the thymus (Magaletta et al., 2020). Impairment of PE formation during PA development was found to be the cause of severe developmental-related abnormalities that are responsible for one-third of all congenital disorders, mainly being tied to a weakened or absent formation of this microenvironment (Jones & Trainor, 2004). Among them, 22q11.2 Deletion Syndrome (22q11.2DS), the most common microdeletion syndrome that affects 1/4000 live births (Baldini, 2006; McDonald-McGinn et al., 2015; Motahari et al., 2019) has been linked to defective PE development. Despite the fundamental role of the PE during PA development and its connection with developmental diseases, the transcriptional and epigenetic dynamics which characterize this cell type remain poorly studied. In order to derive functional PE cells, the *in vitro* differentiation protocols should mimic the sequential origin of intermediate cell types occurring during *in vivo* development (Magaletta et al., 2020). These stages include the specification into Definitive Endoderm (DE), the patterning into Anterior Foregut Endoderm (AFE), and the subsequent specification into PE. Although many groups have worked on the generation of DE and AFE lineages (Green et al., 2011; Kearns et al., 2013; Loh et al., 2014), most of the protocols available so far (reviewed by Magaletta et al., 2020) were only able to generate cells that displayed a moderate level of expression of only a few PE markers (PAX9, SOX2, FOXA2, TBX1) (Green et al., 2011; Kearns et al., 2013) and, in some cases, the cells expressed markers of the DE stage that should have been instead silenced at the PE stage (i.e. SOX17) (Kearns et al., 2013). Furthermore, none of these works ever transcriptionally or epigenetically characterized the cells, leaving a gap in critical information necessary to study this process. Retinoic acid (RA) signaling was shown to be involved in the regulation of the pharyngeal patterning (Koop et al., 2014) and in the proper formation of the third and fourth pharyngeal arches (Kopinke et al., 2006; Wendling et al., 2000). Furthermore, alterations in RA concentration cause defects in the development of the thymus and parathyroids, both structures originating from the 3^rd^ pharyngeal pouches (Begemann et al., 2001; Mulder et al., 1998; Niederreither et al., 2003; Vermot et al., 2003) and complete loss of RA synthesis in the developing embryo recapitulates most of the phenotype of the 22q11.2DS (Vermot et al., 2003). Under the hypothesis that RA might play a crucial and yet under-investigated role in the development of PE cells *in vitro*, we developed and validated a new monolayer differentiation protocol using small molecules in combination with a specific RA concentration and chemically defined media to generate a transcriptionally homogeneous cell population expressing all the PE markers known in literature so far. By combining downstream analysis such as bulk and single-cell RNA-Seq, the Assay for Transposase-Accessible Chromatin with Sequencing (ATAC-Seq), and Chromatin Immunoprecipitation followed by Sequencing (ChIP-Seq) of histone modifications, we were able to deeply characterize the transcriptomic and the epigenomic landscape of our PE cells, to generate a gene regulatory network (GRN), and to identify previously unknown CIS-regulatory elements likely responsible of the proper PE differentiation. In addition, we dissected the transcriptional and epigenetic contribution of RA in PE specification, elucidating in part the role of RA in human pharyngeal development. Our data provides a powerful discovery platform and a valuable resource that can be used for the functional characterization of previously unknown regulatory elements and provides a powerful tool for the study of this important, but still mostly unexplored developmental cell type.

## RESULTS

### Differentiation of hESCs into *bona fide* PE cells by the dynamic exposure of AFE cells to RA signaling and transcriptional characterization

In order to generate a robust and homogenous population of PE cells, we attempted to build upon our previously published protocol for the generation of functional AFE cells (Loh et al., 2014). H9 hESCs (D0) were differentiated into DE (D2) by using the PSC Definitive Endoderm Induction kit, formulated by using the findings from our previous work (Loh et al., 2014), which is now routinely used for the generation of highly pure DE cells. After 24 hours in Medium A and 24 hours in Medium B (**Figure 1A**), the DE cells were anteriorly patterned into AFE by dual inhibition of TGFb and BMP4 for 24 hours (D3). Given the previously described contribution of RA in the PE formation, we tested the hypothesis that the addition of RA was necessary and sufficient to activate a gene regulatory network able to induce the differentiation of AFE into PE-like cells. To test our hypothesis, AFE cells were cultured with increasing concentrations of RA (0-800 nM) and checked for the expression of the known PE markers TBX1, NKX2-5, PAX9, PAX1, FOXA2, and RIPPLY3 (**Figure S1A**). The titration showed that 200 nM represents the optimal concentration, leading to a proper combination of expression of the tested PE markers (**Figure S1A**). Assuming 200 nM as the ideal concentration of RA for the generation of PE, we then decided to identify the optimal exposure time of cells to RA by evaluating the expression of such markers during a 7 days long time course of differentiation and in the presence or in the absence of RA, which was added for 24, 48, 72, or 96 hours (**Figure 1A**). This analysis led us to choose 48 hours as the optimal window of exposure to RA for obtaining PE cells, based on the expression peak of several PE markers (**Figure 1B**). Notably, SOX17, a specific marker of the DE stage, was properly downregulated during the differentiation in both conditions, while the presence of RA reduced the upregulation of PAX1, providing a better correlation with the dynamic expression of this gene observed *in vivo* (Mulder et al., 1998). To move beyond the reductionist approach based on the observation of a few, manually selected marker genes, we decided to deeply characterize and compare the transcriptome of our cells. To do this, polyadenylated RNA from hESCs (D0), DE (D2) AFE, (D5), and PE (D5+RA) cells was collected and submitted to a bulk RNA-Seq analysis. As shown by the sample clustering analyses (**Figure S1B** and **C**), AFE and PE samples have similar but distinct gene expression profiles, significantly different from those of hESCs and DE samples, which form two clearly separated clusters. Differential gene expression analysis performed between each pair of conditions allowed us to identify 7226 differentially expressed genes (DEGs) (**Figure S1D**,). We then focused on the DEGs with great variation in expression (see Methods) among the DE, PE, and AFE conditions, for a total of 4578 genes. Such genes were grouped into ten clusters based on their expression trends (**Figure 1C**). Gene Ontology (GO) term (Ashburner et al., 2000) enrichment analysis was performed on each cluster to gain insight into the function of each class of DEGs **(Figure 1D**,). While we found that genes from all the clusters were involved in Biological Process (BP) categories related to development and morphogenesis, we also observed that the “pharyngeal system development” was highly enriched in cluster 3, which is composed of genes whose expression is induced from DE to AFE-PE and particularly boosted by the presence of RA. In agreement with this functional enrichment analysis, all the known marker genes of the PE are upregulated during the transition from DE to AFE and PE, and most of them are induced by the presence of RA **(Figure 1C**, green boxes), while DE-specific markers are properly downregulated (**Figure 1C**, orange boxes). To further confirm the reliability of our protocol in activating a PE-specific transcriptional network, we identified the functional categories enriched among upregulated and downregulated genes in DE vs AFE and DE vs PE contrasts *via* Gene Set Enrichment Analysis (GSEA) (Subramanian et al., 2005). As expected, for both comparisons we found the “endoderm development” GO BP category to be enriched among the downregulated genes (i.e. genes more expressed in DE) as well as the “pharyngeal system development” and related categories to be over-represented among the upregulated genes (**Figure S1E**, upper panels). Furthermore, by performing GSEA on the AFE vs PE contrast to highlight the major transcriptional changes induced by the addition of RA, we found the “activation of HOX genes during differentiation” and “activation of anterior HOX genes” Reactome pathways (Gillespie et al., 2022) as the most enriched among the upregulated genes (**Figure S1F**). Finally, we compared the expression profile of our cells with that of the mouse *in vivo* counterparts by taking advantage of the single-cell transcriptomic data previously published by Han and colleagues (Han et al., 2020), who generated an elegant spatiotemporal map of endoderm and mesoderm development during murine foregut organogenesis (**Figure S1G**). Interestingly, we found HOXA1, HOXA2 HOXB1, and HOXB2 genes, the most definitive regulators of Anterior-Posterior patterning, to be specifically upregulated in our PE cells and in the Pharyngeal endoderm clusters of mouse *in vivo* development (e_b3 and e_c3) (**Figure S1G**). These results demonstrate that RA signaling broadly regulates the Anterior-Posterior HOX code during patterning of the human gut tube, which likely leads to the activation of a proper transcriptional network necessary to generate PE cells. Finally, we compared our cells with the different endodermal clusters identified by Han and colleagues (**Figure 1E**) by looking at the transcription factor (TF) expression profile. Notably, we found AFE cells to be similar to the corresponding anterior foregut cluster in mouse **(Figure 1E**, e_a5), while PE cells showed high similarity with the Pharyngeal Endoderm clusters at day 9.0 (**Figure 1C**, e_b3 and e_b4) and at day 9.5 (**Figure 1C**, e_c3), further confirming that, based on their gene expression profile, these cells can be considered *bona fide* human Pharyngeal Endodermal cells.

**Figure 1:**
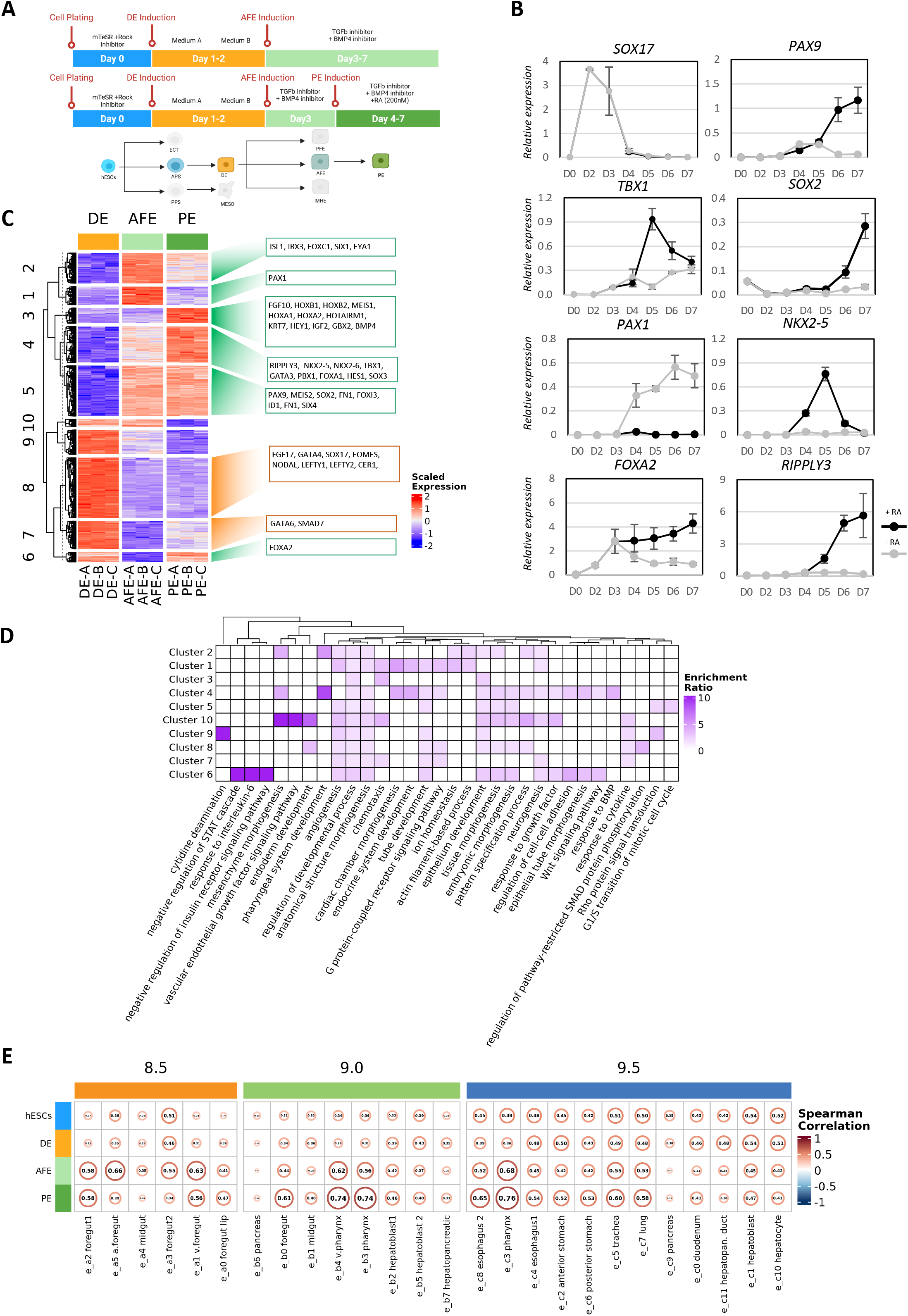
Transcriptomic analysis of *in vitro* derived pharyngeal endoerm cells. **(A)** Schematic representation of the protocol for the *in vitro* differentiation of hESCs into Pharyngeal Endoderm cells. Created with BioRender.com. **(B)** RT-qPCR time course analysis showing the relative expression of DE and PE specific markers in the presence (black) or absence (grey) of RA. Data were normalized on PDGB expression and represent means ± SEM of three independent time-course experiments. **(C)** Heatmap showing the expression in DE, AFE and PE samples of the greatly varying DEGs identified in DE vs AFE and DE vs PE contrasts, as well as their separation in ten clusters produced *via* k-means clustering. Hierarchical clustering of the ten clusters is also shown. Known markers belonging to each cluster are shown in the boxes on the right. The expression values reported in the heatmap correspond to row-scaled (Z-score), rlog-transformed count data. **(D)** Heatmap showing the results of the GO BP term enrichment analysis performed on the ten gene clusters shown in C; the color intensity in each cell is proportional to the Enrichment Ratio. The heatmap reports only a set of the significantly enriched categories (FDR < 0.05 in at least one cluster), selected in order to reduce redundancy; Enrichment Ratio is plotted only when FDR < 0.25. **(E)** Spearman correlation matrices showing the similarity between hESCs, DE AFE, and PE cells and embryonic mouse foregut endodermal cell type clusters identified in Han et al., 2020, based on the expression of the top 10 transcription factors enriched in each cluster. Text and circle width in each cell of the matrices are proportional to the absolute value of the Spearman correlation. Each matrix corresponds to a different murine developmental stage (day 8.5, day 9.0 and day 9.5).

### scRNA-Seq analysis reveals a homogeneous transcriptional signature for both DE and PE differentiated cells

To comprehensively decipher the transcriptional dynamics occurring during the differentiation into PE at the single-cell level, we performed single-cell RNA Sequencing (scRNA-Seq) of hESC, DE, and PE samples, with a yield of 6115, 2566, and 2525 cells, respectively. A total of 4887, 1127, and 1115 cells passed the quality controls (see Methods for further details). The expression of highly variable genes was used to visualize cells in two dimensions with the Uniform Manifold Approximation and Projection (UMAP) method (Becht et al., 2019), which shows a clear separation of cells from the three different stages (**Figure 2A**, top panel).

**Figure 2:**
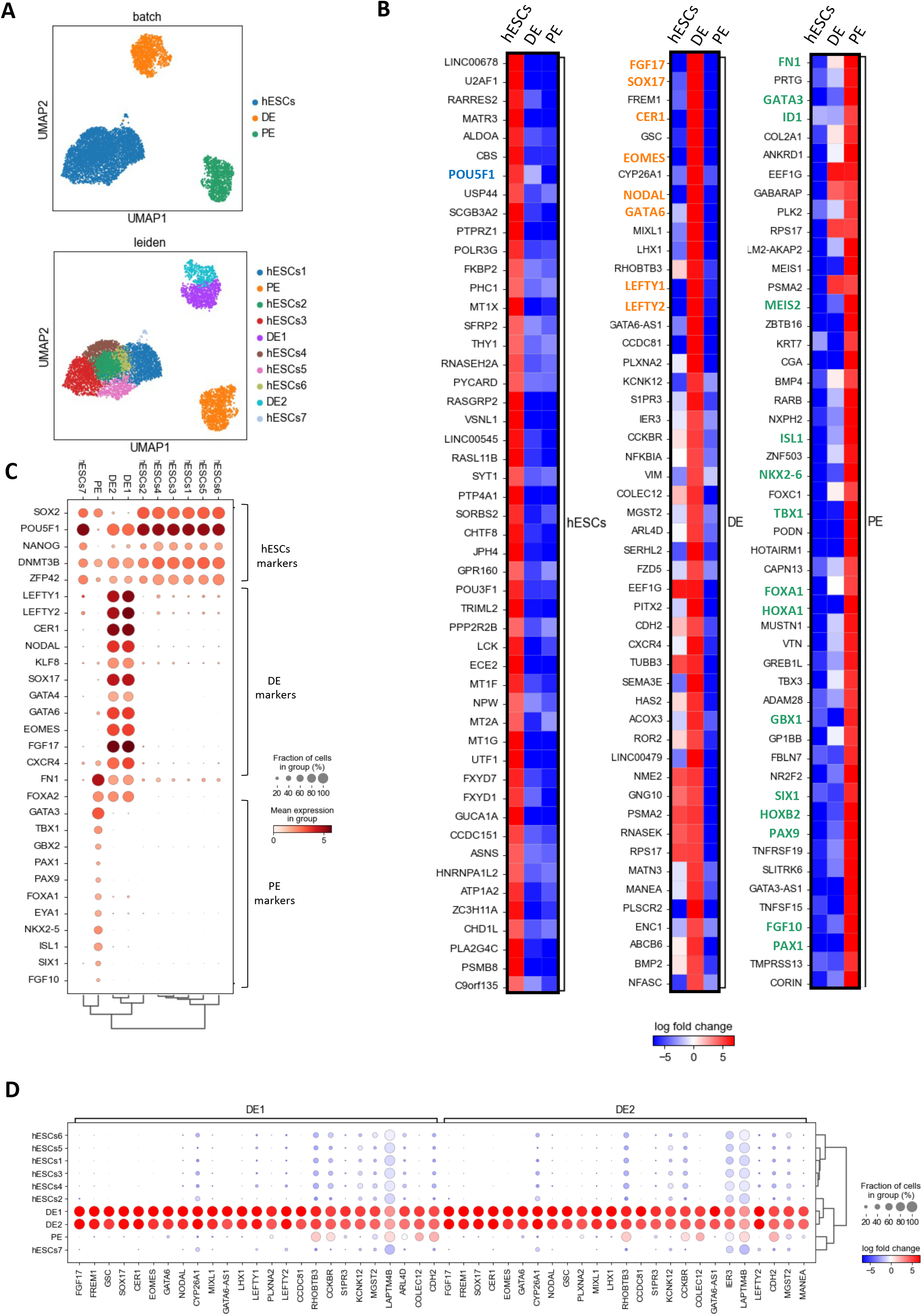
scRNA-Seq data reveals the homogeneity of the *in vitro*-derived DE and PE cells. **(A)** UMAP-based visualization of scRNA-Seq data produced from hESCs, DE and PE cells, with cells colored based on the differentiation stage (top panel) and Leiden clustering (bottom panel). **(B)** Heatmaps showing the log2(FC) of top 50 DEGs identified in the hESCs, DE and PE clusters (log2[FC] >= 4). Known markers for the hESCs, DE and PE stages are highlighted in blue, orange and green, respectively. **(C)** Dot plot showing the expression (log[norm.counts+1]) of known hESCs, DE and PE markers in each of the 10 different clusters. The dot size represents the percentage of cluster cells expressing the gene and the color represents the average expression in the cluster. The clustering tree is shown at the bottom of the chart. (**D**) Dot plot of the top 25 genes responsible for the DE clusters identity (x-axis) and their expression (log[norm.counts+1]) in the 10 different clusters (y-axis). The dot size represents the percentage of cluster cells expressing the gene and the color represents the log2(FC) of the gene expression compared to the other clusters.

We subsequently identified the top 50 DEGs for each time point; this analysis confirmed the upregulation of known hESCs, DE, and PE markers respective to their specific cell stage (**Figure 2B**), further supporting the robustness of our protocol. Next, we investigated the transcriptional heterogeneity of our cells by applying the Leiden clustering algorithm (Traag et al., 2019). This analysis identified 10 different clusters based on their transcriptomic similarities (**Figure 2A**, bottom panel). Interestingly, all the cells belonging to the PE group were assigned to a single cluster (PE), supporting our hypothesis that our *in vitro* differentiation protocol leads to the formation of a transcriptionally homogeneous cell population, while hESCs and DE cells were divided into 7 and 2 clusters, respectively.

Next, we evaluated the expression of known hESCs, DE, and PE marker genes within each cluster to understand whether they correspond to cells having a significantly different biological identity within the same cell population (**Figure 2C** and **Figure S2A**). As shown in **Figure 2C**, all the hESCs clusters uniformly express hESCs markers such as POUF51, SOX2, NANOG, DNMT3B, and ZFP42; the same situation was observed for DE clusters, which are homogeneously positive for known DE markers such as LEFTY1, LEFTY2, CER1, KLF8, SOX17, GATA4, GATA6, EOMES, and CXCR4. As expected the PE cluster expresses known PE markers such as FN1, GATA3, TBX1, PAX1, PAX9, EYA1, NKX2-5, ISL1, and SIX1, while FOXA2 expression was detected in both DE and PE cells, and SOX2 expression resulted to be particularly evident in hESCs and PE cells. According to this analysis, hESC1-7 and DE 1-2 clusters express the expected marker genes despite their assignment to different clusters. We also evaluated the expression of the top genes responsible for the identity of each cluster (**Figure 2D** and **Figure S2B**). The nearly identical expression profile of such genes among clusters of the same differentiation stage, along with the great overlap of these genes, confirmed the transcriptional similarity of the clusters identified *via* the Leiden approach for each stage. This conclusion was further supported by the fact that most of the 50 top DEGs identified in the DE1 vs DE2 contrast are not stage-specific, since they are also expressed in PE and hESCs stages (**Figure S2C**). Taken together, these data suggest that both PE and DE represent a homogenous cell population that properly differentiates into the expected cell type.

### Retinoic Acid induces chromatin accessibility changes accompanying transcriptomic variations during PE differentiation

To provide a clear picture of the putative regulatory regions responsible for the activation of a PE-specific transcriptional program, in particular in response to RA signaling, we decided to deeply characterize and investigate the epigenetic landscape of DE, AFE, and PE cells *via* ATAC-Seq. Peak calling was performed for each sample to find accessible regions; consensus peaks for each condition and peaks in common between different conditions were subsequently identified. Following this approach, we discovered 107569 accessible regions in DE, 95231 in AFE, and 80441 in PE, 50015 of which are in common among the three conditions (**Figure S3A**, upper panel). Clustering analyses performed on ATAC-Seq samples clearly showed that our cells have a distinct epigenomic landscape at each stage, the AFE and PE cells being more similar to each other than to DE cells as observed from transcriptomic data, confirming the quality and the reproducibility of each replicate (**Figure S3B** and **S3C**).

To locate the genomic regions responsible for each stage-specific epigenetic profile, we performed a differential accessibility analysis between each pair of conditions, which led to the identification of differentially accessible regions (DARs). For each comparison, DARs were classified into Gain (log2 fold change [FC] > 1) and Lose (log2[FC] < −1) peaks (**Figure S3A**, lower panel). Based on read coverage, DARs were further grouped into six different clusters (**Figure 3A**); each of them shows a distinct behavior during differentiation, indicating that the chromatin accessibility is actively changing during the induction of the differentiation and that is actively responding to the addition of RA. Concordantly with the number of consensus peaks observed in each condition, the largest cluster is the one composed of regions whose accessibility decreases in the transition from DE to AFE and PE (cluster 1, 30927 peaks), indicating that a significant proportion of genomic regions are closed during differentiation. Interestingly, an analysis of the evolutionary conservation in vertebrates performed on DARs from each cluster showed that, overall, DARs tend to be more conserved than regions with no significant change in accessibility (Common peaks). Moreover, regions gaining accessibility in response to differentiation and regions specifically open or closed upon the addition of RA (clusters 3, 4 and 5) have higher levels of sequence conservation than other DARs, suggesting an evolutionarily conserved function for such sequences (**Figure 3A**) and that the mechanism through which RA promotes the transcriptional and epigenetic maturation of PE could be similar in other species. Another feature that distinguishes DARs from Common peaks is their genomic distribution with respect to gene elements (**Figure 3B**): while ∼33% of Common peaks are in promoter regions, DARs from all clusters are more often located outside such regions. This is particularly evident for peaks losing accessibility in the transition from DE to PE (cluster 1 and cluster 3, ∼9% of the peaks falling in promoter regions), and less so for DARs specifically open in both AFE and PE (cluster 5) and only in PE (cluster 4) (∼17% and ∼14% of the peaks falling in promoter regions, respectively). DARs not overlapping with promoter regions could regulate the expression of nearby genes by acting as enhancers. To test this hypothesis, for each DAR cluster we identified a gene set composed of genes whose transcription start sites (TSSs) are less than 50 kilobases (kb) away from any of the peaks of the cluster; then, for each differential gene expression contrast, we performed a GSEA to evaluate whether cluster-specific gene sets are enriched among the upregulated or the downregulated genes (**Figure 3C**). The results of this analysis showed that Gain DARs tend to be located near upregulated genes, while Lose DARs are found in the proximity of downregulated genes. The functional significance of DARs, inferred by the function of their nearby genes, was investigated by performing GREAT analysis (McLean et al., 2010) on each peak cluster (**Figure 3D**). Interestingly, we found a clear functional enrichment for Clusters 5 and 4 in GO BP terms related to the development of the Pharyngeal Apparatus (with Cluster 5 showing enrichment for genes involved in thymus development, a downstream cell type originating from the PE), further supporting the notion that chromatin is dynamically inducing the establishment of a transcriptional program promoting the differentiation of our cells into PE progenitors. To gain insight into the contribution of TSS accessibility changes to shaping the observed variations of gene expression, we evaluated the ATAC-Seq read coverage around the TSS of previously identified DEGs and of non-DEGs. Interestingly, we observed a general increase in the TSS accessibility in the transition from DE to AFE and PE both in DEGs and non-DEGs (**Figure S3D**), and some enrichment of Gain and Lose peaks in the promoters of upregulated and downregulated genes, respectively (**Figure S3E** and **F**). However, since most of the DEG TSSs overlap with Common ATAC-Seq peaks (**Figure S3E**), it appears that dynamic promoter accessibility is not a dominant effect in the regulation of gene expression and that the chromatin accessibility changes responsible for the variations in gene expression mainly occur in regions located outside promoters. In support of this, we found several DE and PE markers whose nearby regions are respectively closing and opening during the differentiation (**Figure S3G**).

**Figure 3:**
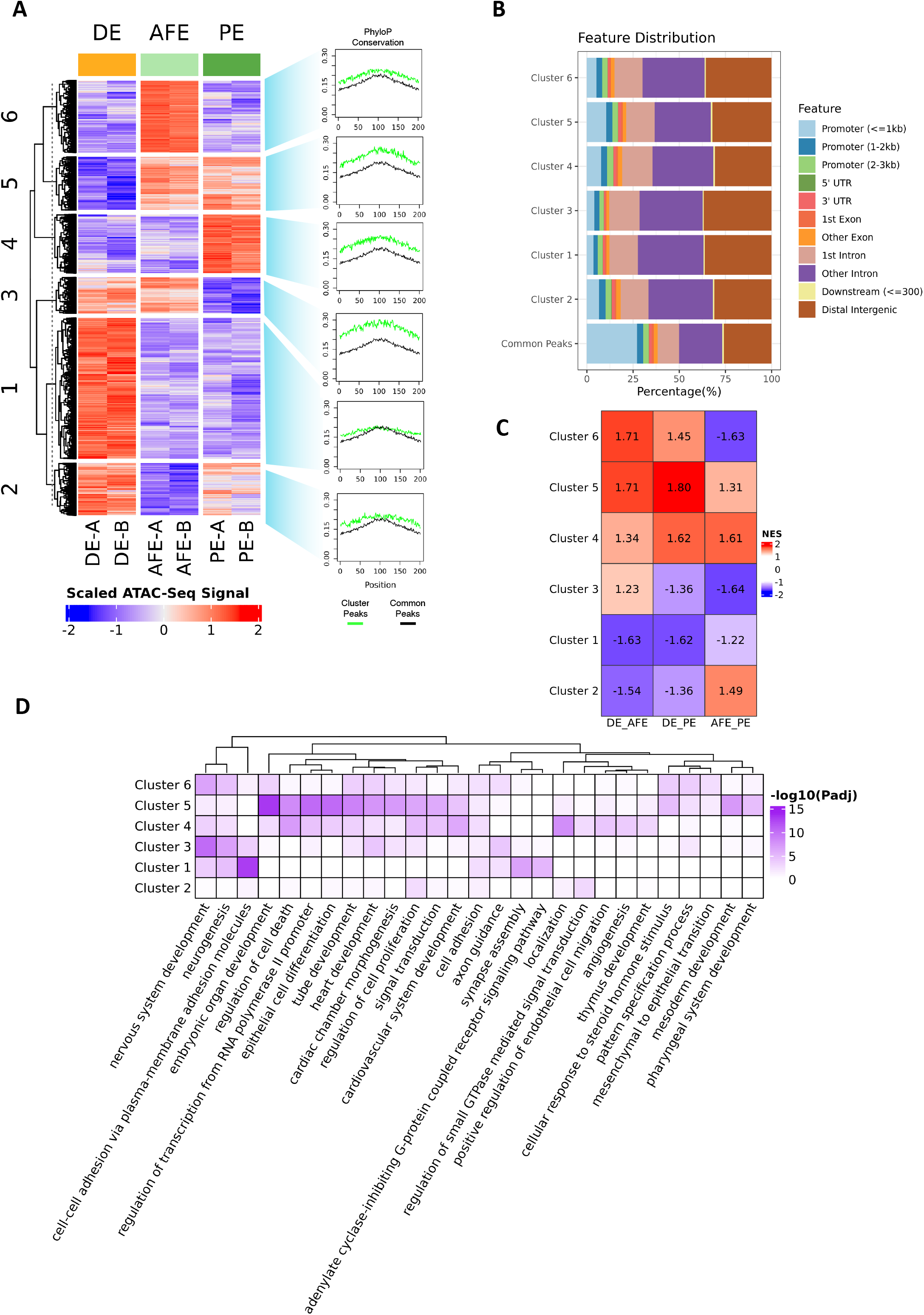
ATAC-Seq analysis of DE, AFE and PE cells. **(A)** Heatmap showing the ATAC-Seq signal in DE, AFE and PE samples of the DARs identified in all contrasts, as well as their separation in six clusters produced *via* k-means clustering. Hierarchical clustering of the six clusters is also shown. Average PhyloP conservation scores, calculated for each genomic position within DARs and Common peaks, are shown in the plots on the right. The ATAC-Seq signal values reported in the heatmap correspond to row-scaled (Z-score.) log2-transformed library size-normalized count data. **(B)** Bar plot showing the genomic annotation of DARs belonging to each cluster shown in Figure 3A and Common ATAC-Seq peaks. Each genomic feature is represented by a specific color shown in the legend. **(C)** Table showing the Normalized Enrichment Scores (NES) calculated performing GSEA on each differential gene expression contrast (DE vs AFE, DE vs PE and AFE vs PE) and using sets of expressed protein-coding genes having a TSS in proximity (< 50 kb) of cluster-specific DARs. Positive NES: the gene set is enriched among the upregulated genes; Negative NES: the gene set is enriched among the downregulated genes. **(D)** Heatmap showing the results of the GREAT analysis performed on the six DAR clusters shown in A; the color intensity in each cell is proportional to the adjusted p-value. The heatmap reports a set of the significantly enriched GO BP terms (adjusted p-value < 0.01 in at least one cluster), selected in order to reduce redundancy.

### Prediction of stage-specific transcription factor activity points to a major role of RA in PE differentiation

The significant sequence conservation of the DARs we identified in the transition from DE to AFE and PE (**Figure 3A**) suggests that such regions might be involved in regulating the differentiation process, possibly *via* the binding with protein regulators. Given the well-known role of transcription factors (TFs) in establishing transcriptional networks responsible for proper cell differentiation, we decided to investigate the putative TF binding profile of DARs. To this end, we used the maelstrom tool (Bruse & van Heeringen, 2018) to perform a differential motif enrichment analysis revealing which known TF motifs are specifically enriched in cell type-specific accessible regions. In parallel, the BiFET tool (Youn et al., 2019) was employed to identify TF footprints (FP: less accessible regions within highly accessible regions where a TF motif is found) (Li et al., 2009) enriched in the DARs found in each differential accessibility contrast (). The results of the maelstrom and of the BiFET analyses were integrated by selecting the TF motifs whose differential enrichment trend correlates with the corresponding FP enrichment profile and with the TF expression during the differentiation. This way we identified a robust set of transcription factors, most of which are known regulators of DE, AFE or PE differentiation, which are differentially active among the three cell types (**Figure 4A**). Notably, among the TFs with PE-specific activity, we found several known regulators of PE differentiation such as FOXA1, FOXA2, NKX2-5, GATA3, PAX9, MEIS1, MEIS2, HOXA1 and HOXB1/2 (**Figure 4A**), supporting the idea that RA regulates the accessibility of chromatin regions that are functionally relevant to PE commitment. As expected, among the motifs enriched in regions gaining accessibility in PE, we also found two DR5 type Retinoic Acid Responsive Elements (RAREs), whose enrichment correlates with the expression of RARA and RARB, two Retinoic Acid receptors (RARs) that are activated after the binding with RA and mediate the cellular response to this morphogen.Gain DARs harboring enriched RARE FPs (also including DR1 and DR2 RAREs) are mostly distributed in non-promoter regions, showing the same trend of Gain DARs without RAREs (**Figure S4A**). We then investigated whether such Gain regions putatively bound by RARs could influence the expression of nearby genes. To do this, we first identified Gain DARs with a RARE FP located in promoters or non-promoter regions and evaluated the expression status of genes regulated by those promoters or genes harboring the closest TSSs (< 50 kb), respectively, as compared to that of genes found in the proximity of Common peaks (**Figure 4B**). Interestingly, we found a significant enrichment in AFE vs PE upregulated genes only in the proximity of Gain peaks with RAREs located in non-promoter regions, suggesting that the RA might regulate the expression of those genes by interacting with intergenic regions with enhancer activity. By performing a GO BP term enrichment analysis on such genes, we found a significant enrichment for developmental categories such as “tube development” and “animal organ morphogenesis” (**Figure 4C**), which are processes known to be regulated by the presence of RA. Interestingly, among the upregulated genes belonging to the enriched categories, we found HOXA1 and HOXB1, known RA targets. By looking at genomic regions spanning the HOXA1/2 and HOXB1/2 gene loci, we found several spots with increased accessibility, many of which harboring a RARE FP (**Figure S4B**). Notably, among them we found three Gain DARs, one proximal to HOXA1 and two close to HOXB1, which correspond to RA-responsive enhancers that are known to activate the expression of these genes both in mouse and human (Huang et al., 2002; Langston et al., 1997; Marshall et al., 1994; Ogura & Evans, 1995; Thomson et al., 1998).

**Figure 4:**
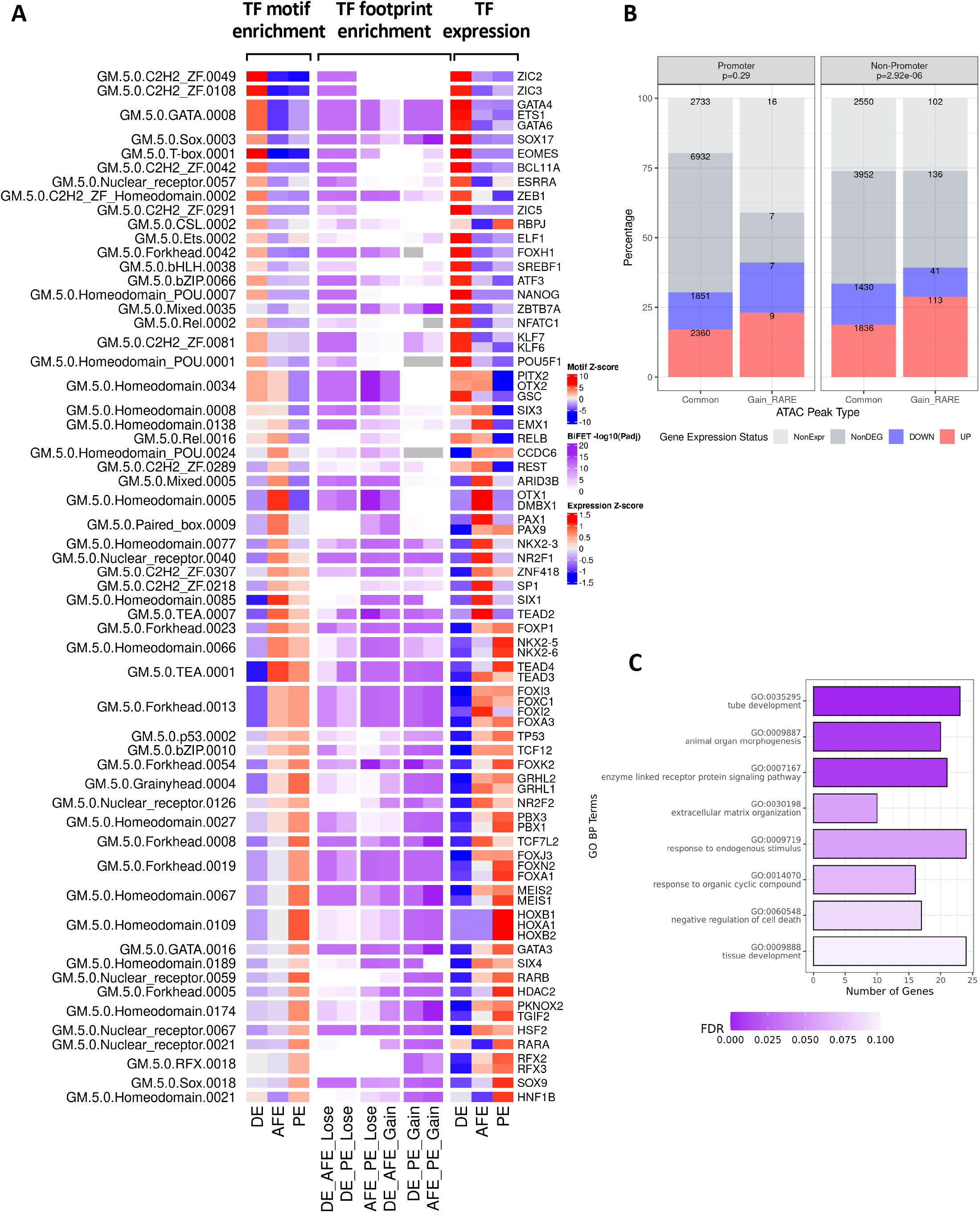
TF binding prediction on DE, AFE and PE cells. **(A)** Heatmaps showing the results of the integrated analysis of cell type-specific TF activity. The TF motifs here reported were selected based on maelstrom Z-score (heatmap on the left), BiFET adjusted p-value (heatmap in the middle) and TF expression (heatmap on the right), and on the correlation between these measures (see Methods). The motifs spanning multiple rows are associated to multiple TFs having expression correlated with enrichment. **(B)** Bar plot showing, for the AFE vs PE contrast, the expression status of the protein-coding genes whose promoter hosts Common ATAC-Seq peaks or Gain peaks with RARE FP elements (left panel) and of the protein-coding genes closest to the non-promoter Common peaks or Gain peaks with RARE FP elements (right panel). For each non-promoter peak, the nearest gene was identified by searching for the closest TSS of an expressed gene within 50 kb; if no expressed gene was found among the three closest ones, the nearest non-expressed gene within 50 kb was chosen. The p-values reported in each plot were calculated by comparing the proportions of upregulated genes between the two classes of genes *via* Fisher’s exact test. NonExpr: average gene TPM < 1 in both AFE and PE); NonDEG: the gene is not differentially expressed; DOWN: the gene is downregulated in PE (log2[FC] significantly < 0); UP: the gene is upregulated in PE (log2[FC] significantly > 0). **(C)** Bar plot showing the GO BP terms enriched (FDR < 0.1) among the upregulated genes identified in the AFE vs PE contrast that are in proximity of Gain ATAC-Seq peaks hosting RARE FP elements located in non-promoter regions. The X-axis reports the number of such genes that belong to each enriched category; the color intensity is inversely proportional to the enrichment FDR.

### Epigenetic characterization of regulatory elements reveals functional chromatin state changes between DE and PE stages

To further define and complement the epigenetic landscape of DE to PE differentiation, we performed ChIP-Seq experiments of H3K4me_3_, H3K27me_3_, H3K4me_1_, and H3K27ac histone marks (HM) on chromatin isolated from DE and PE cells. In addition to showing a clear agreement between biological replicates, the hierarchical clustering of samples based on ChIP-Seq read coverage also revealed that the two cell types have distinct HM profiles (**Figure S5A**). The proper HM distribution was confirmed by evaluating the DE and PE HM depositions around (+/- 3 kb) the TSSs of protein-coding genes, stratified based on the presence (or absence) of ATAC-Seq peaks, and around the summits of ATAC-Seq peaks located outside promoter regions (**Figure S5B** and **C**). H3K4me_3_ and H3K27ac show a bimodal distribution centered on the TSS, with a greater occupancy at sites where the chromatin is open both in DE and in PE (**Figure S5B**); differentially accessible TSSs display a clear and concordant change in the H3K27ac signal. Similarly, the H3K27me_3_ and H3K4me_3_ deposition at TSSs depends on chromatin accessibility dynamics, with a clear drop in the signal that is evident only at open TSS sites with no significant change in accessibility between DE and PE (**Figure S5B**). As expected, the HMs that were predominantly found within non-promoter ATAC-Seq peaks were H3K4me_1_ and H3K27ac, whose deposition positively correlates with the differential chromatin accessibility between DE and PE, concordantly with their well-established role as markers of regions with enhancer activity (Andersson & Sandelin, 2020) (**Figure S5C**).

Taking advantage of the well-known distinct epigenetic signature of different functional genomic elements (Andersson & Sandelin, 2020), we annotated the epigenome of DE and PE cells based on the presence of specific HM combinations (chromatin states) using the ChromHMM software (Ernst & Kellis, 2012). We selected a 10-state model as the one which better and more concisely describes the meaningful combinations between the HMs under investigation (**Figure 5A**); the human genome was segmented into 200 bp bins and each of such intervals was annotated with the states found in DE and PE. Based on the function that is commonly associated to known HM combinations (van der Velde et al., 2021) and on their overlap with annotated functional regions (**Figure 5B**), we renamed the model states to: TssA (active/acetylated Promoter), Tss (Promoter), TssFlnk (Tss flanking region), TssBiv (bivalent promoter), ReprPC (Polycomb-repressed), EnhA (active/acetylated enhancer), EnhPr (primed enhancer), EnhBiv (bivalent enhancer), Quies1 and Quies2 (quiescent regions with no histone marks, fused into Quies state in subsequent analyses). Looking at how the genomic distribution of functional chromatin states changes in the transition from DE to PE, we observed a clear decrease in the number of genomic regions repressed by Polycomb and occupied by active enhancers, and an increase in primed enhancer occupancy (**Figure S5D**). Interestingly, by evaluating the overlap between chromatin states and ATAC-Seq peaks, we found that, while a significant amount of Common peaks is located within regions with promoter-specific histone marks combinations, DARs are less frequently found in such regions and are enriched in enhancer-related signatures (**Figure 5B** and **S5E**), in line with the previously discussed genomic distribution of accessible regions. Furthermore, DARs are more frequently found in a quiescent state in the cell type in which the chromatin is less accessible (**Figure S5E**), indicating that, as expected, chromatin opening and closing events are accompanied by a concomitant change in histone mark deposition. To note, we also found that DE bivalent promoters are enriched in Gain peaks, suggesting that the activation of genes controlled by these promoters in the DE-PE transition might also be the result of an increase in chromatin accessibility. We then sought to characterize the chromatin state change dynamics in the transition between the DE and PE stages. To this end, for each transition between two distinct chromatin states observed in the differentiation process, we evaluated its enrichment with respect to the same change in the opposite direction (see Methods) (Fiziev et al., 2017) (**Figure 5C**). The most relevant state changes emerging from this analysis are those going from bivalent or Polycomb-repressed states in DE to active and primed promoters and enhancers in PE. This is in line with the well-known gradual resolution of bivalent chromatin domains (Bernstein et al., 2006) during cell differentiation. We also observed a strong transition from active enhancers to quiescent states, which is concordant with the high number of intergenic regions losing accessibility in the differentiation from DE to PE, and a significant shift from deacetylated to acetylated promoters (**Figure 5C**). To further validate the robustness of our model and to identify functional genomic regions associated to changes in gene expression and chromatin accessibility, we compared the state transitions with the expression of nearby genes (**Figure 5D**) and evaluated the dynamics of overlapping ATAC-Seq peaks (**Figure 5E**). As expected, those genes which are found in the proximity of genomic regions transitioning from an active to a repressed state tend to be downregulated, while transitions from repressive and quiescent states to active states are enriched in upregulated nearby genes (**Figure 5D**). The state transitions showing the greatest overlap with Gain DARs are those leading to the formation of active and primed enhancers, while the regions in which such elements are lost are enriched in Lose DARs (**Figure 5E**). Notably, the transition to a primed enhancer state displayed a strong association with an increase in nearby gene expression and chromatin accessibility, in line with a recent report showing that enhancers can activate the expression of nearby genes also in absence of the H3K27ac mark in mESCs (Zhang et al., 2020). This effect on nearby gene expression was not observed when the transition to primed enhancers started from quiescent chromatin, except for those cases in which it was accompanied by an increase in chromatin accessibility (**Figure S5F**). We also observed a positive correlation between increased accessibility and transition from a bivalent state to an active promoter (**Figure 5E**). The genes controlled by such promoters showed an enrichment towards development-related functional categories, especially those whose TSS overlaps with a Gain peak, which displayed a specific involvement in the pharyngeal system development (**Figure 5F**). This evidence strongly supports our model and suggests that, during PE differentiation, the chromatin transitions by silencing or selecting regulatory elements, most of which with enhancer signatures.

**Figure 5:**
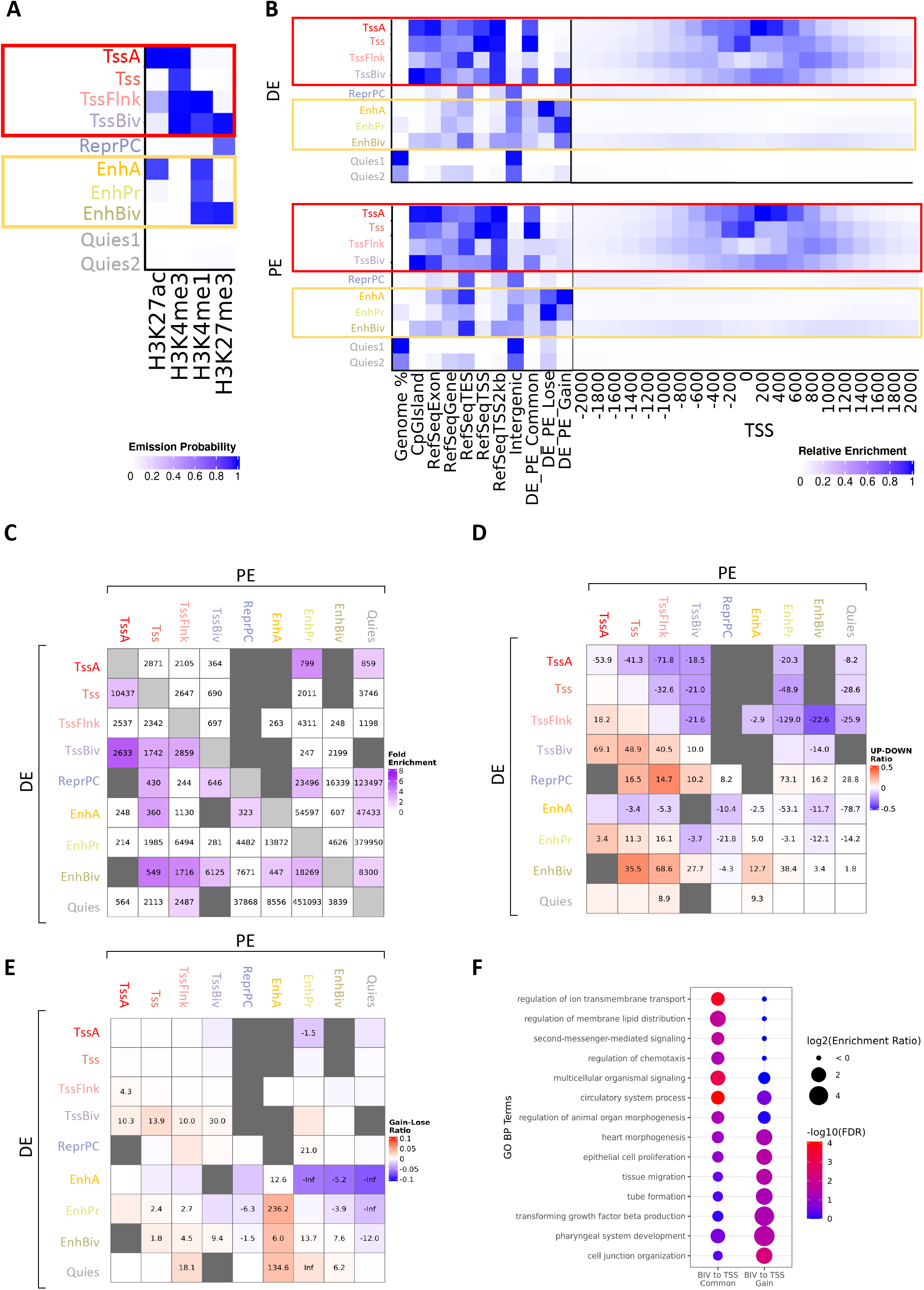
Epigenetic characterization of regulatory elements reveals functional chromatin state changes between DE and PE stages. **(A)** Emission probabilities of the 10-state ChromHMM model. Each row represents a chromatin state and reports the frequency of occurrence of each HM in that state. Red and orange boxes indicate promoter and enhancer states, respectively. **(B)** Heatmaps showing the fold enrichment of each ChromHMM state for different genomic features (left panel) and at fixed positions relative to TSS (right panel) in DE and PE cells. The fold enrichments are calculated as the ratio between observed and expected number of genomic bins for each overlap, except for the Genome % column, which reports the percentage of genomic bins occupied by each state. The color intensities in the left panel are normalized within each column between its minimum value (white) and its maximum value (blue), while those in the right panel are normalized between the minimum value (white) and the maximum value (blue) of the whole matrix. Red and orange boxes indicate promoter and enhancer states, respectively. **(C)** Heatmap showing how many genomic bins transition from a chromatin state to another in the differentiation from DE to PE, as well as the fold enrichment of each transition (see Methods). Only cells corresponding to transitions with fold enrichment > 1.5 are colored, the color intensity being proportional to the fold enrichment. Poorly represented transitions (< 200 bins) are masked using dark grey color. **(D)** Heatmap showing the chromatin state transitions that are enriched in upregulated (red) or downregulated (blue) nearest genes. The color intensity is proportional to the difference between the number of upregulated and downregulated nearest genes, divided by the total number of nearest genes, while the digits within each cell correspond to the –log10(adjusted p-value) of the enrichment, with a – sign when the enrichment is towards downregulated genes. **(E)** Heatmap showing the chromatin state transitions that are enriched in Gain (red) or Lose (blue) ATAC-Seq peaks. The color intensity is proportional to the difference between the number of Gain and Lose overlapping peaks, divided by the total number of bins involved in the transition, while the digits within each cell correspond to the – log10(adjusted p-value) of the enrichment, with a – sign when the enrichment is towards Lose peaks. **(F)** Dot plot showing the GO BP terms enriched among the genes whose promoter transition from a bivalent state in DE (TssBiv, EnhBiv) to a TSS state in PE (TssA, Tss, TssFlnk) and overlap with a Common or Gain ATAC-Seq peak. The heatmap reports only a set of the significantly enriched categories (FDR< 0.05 in at least one class of TSS), selected in order to reduce redundancy.

### Prediction of a transcription factor regulatory network guiding the PE differentiation

To further elucidate the role of TFs in driving the PE development *via* binding to differentially accessible DNA sequences, we performed a FP enrichment analysis on both Gain and Lose DARs after stratifying them based on the overlap with different state transitions (**Figure S6A**). In addition to confirming the importance of the differentially active regulators reported in **Figure 4A**, this analysis also highlighted some differences in the TF binding profile between enhancer and promoter regions – e.g. the TSSs that become active and more accessible in PE are almost exclusively enriched in basic Helix-Loop-Helix (bHLH) TF binding (**Figure S6A**).

In order to identify the key TFs in PE stage determination, we used the ANANSE tool (Xu et al., 2021), which exploits RNA-Seq, ATAC-Seq, H3K27ac ChIP-Seq, and known TF motifs to build a differential enhancer-based network between two different cell types. This differential network was used to identify the set of transcription factors having the greatest influence on the transcriptional changes leading to PE specification (**Figure S6B**). Among them, we found TBX1 and HES1, two TFs active in PE (Arnold et al., 2006; Jackson et al., 2014; Kameda et al., 2013) that did not emerge from our previous enrichment analyses. Finally, we integrated the ANANSE differential network with the TF footprint analysis to infer a GRN which could explain how TFs activate each other by binding to cis-regulatory elements with increased accessibility during PE differentiation. (**Figure 6**). Interestingly, this TF-TF network exhibits a hierarchical structure, in which three classes of TFs can be identified:

- Some TFs, arranged in the upper part of the network, are almost exclusively engaged in outgoing interactions. Many of these TFs, such as FOXA2, KLF3, OVOL2, PAX9, ZNF219, HES4, MEIS1, and TP53, are known master regulators of PE differentiation (Magaletta et al., 2020);
- Other TFs, such as FOXA1, FOXP1, GATA3, HES1, MEIS2, NKX2-5, and RARA, are both activated by the previous ones and involved in the activation of downstream effectors;
- A final set is composed of TFs like HOXA1, HOXB1/2, TBX1, NKX2-6, and PKNOX2, which only have incoming interactions, suggesting that these are the most downstream effectors of the TF-TF activation network.

**Figure 6:**
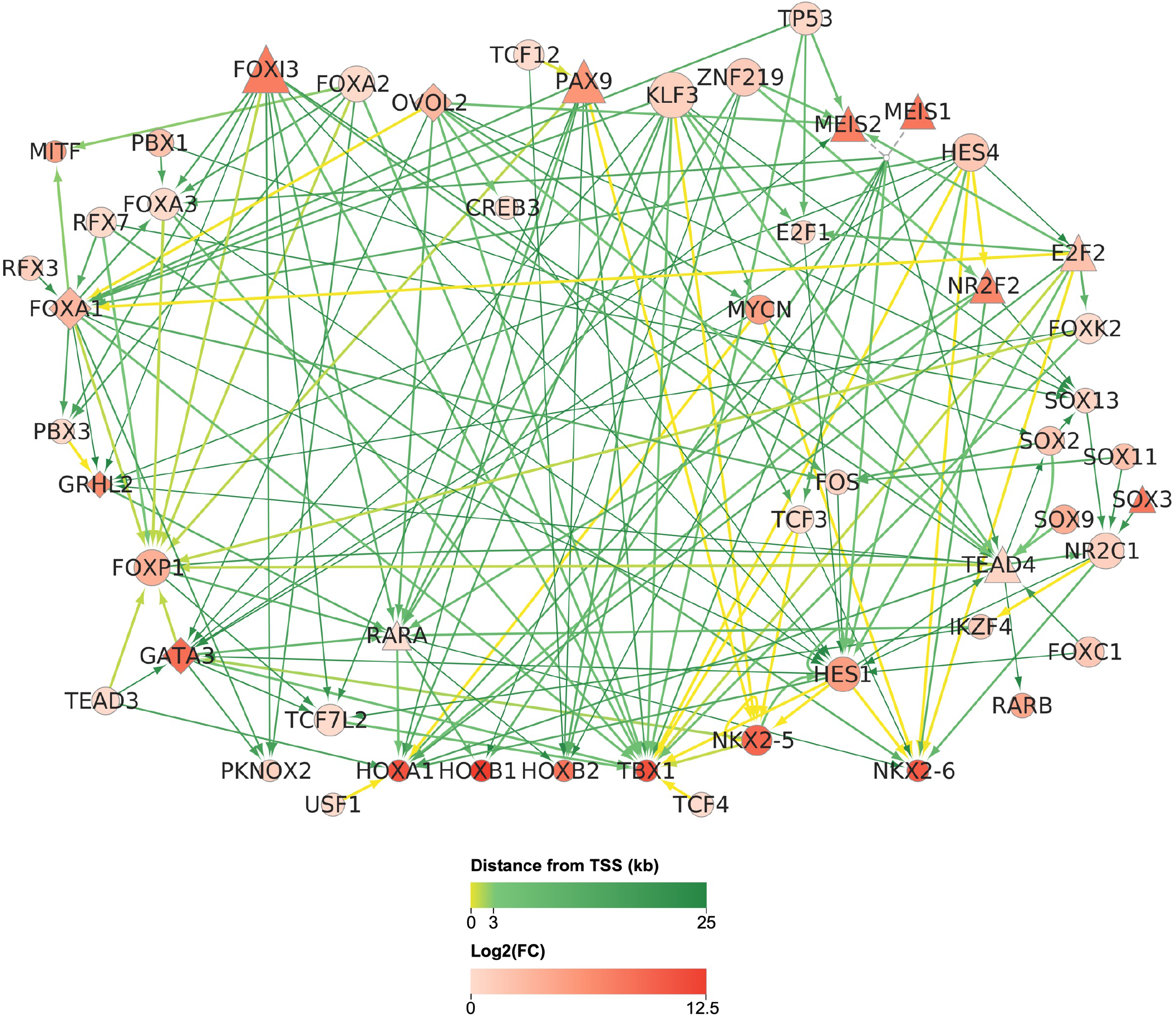
PE Gene Regulatory Network. PE-specific TF-TF activation network. Nodes represent TFs that are specifically active and upregulated in PE. Directed edges indicate the presence of a Gain peak harbouring a source TF-specific FP located less than 25 kb from the TSS of the target TF. Node size is proportional to the number of outgoing interactions. Node color intensity is proportional to the log2(FC) in the DE vs PE contrast. Edge width is proportional to the difference of the ANANSE interaction score between PE and DE (only the edges with differential scores > 0.7 are shown). Edge color represents the minimum distance between the target TSS and a Gain peak with a source-specific FP. TFs whose promoter transitions from a bivalent state in DE (TssBiv, EnhBiv) to a TSS state in PE (TssA, Tss, TssFlnk) are represented with a triangle or a diamond if the transition overlaps with a Common or Gain ATAC-Seq peak, respectively.

## DISCUSSION

The Pharyngeal Endoderm represents a clinically relevant cell type, given its ability to differentiate into organs and apparatuses whose proper formation is affected in several classes of human developmental syndromes displaying a complex spectrum of different phenotypes (Graham et al., 2005, McDonald-McGinn et al., 2015). Revealing the dynamic of human PE cell differentiation and identifying the main players driving this process are essential steps to understanding the pathogenesis of such diseases and to developing effective therapeutic strategies (Magaletta et al., 2020). Nonetheless, the molecular mechanisms underlying human PE development are still mostly unexplored and unknown. Cis-regulatory elements play a pivotal role during cell differentiation, given their ability to interact with tissue-specific TFs and orchestrate spatial and temporal gene expression. Sequence changes within these regions can significantly impact development by altering tissue-specific expression and causing phenotypic variation and diseases (Lidral et al., 2015; Mathelier et al., 2015). In this work, we have filled in part this important knowledge gap by developing a robust, reproducible *in vitro* platform for the derivation of homogenous human PE cells from hESCs through a multi-step protocol involving the addition of known signaling agonists including RA. We have demonstrated that RA is necessary and sufficient to induce the activation of a PE-specific transcriptional network via the remodeling of chromatin structure. Our PE cells have been deeply characterized and profiled based on their epigenetic and transcriptional signatures and appear to be transcriptionally homogeneous and resemble their mouse *in vivo* counterparts. By combining transcriptomic, chromatin accessibility, and histone marks analysis we have outlined the gene regulation dynamics underlying PE differentiation and identified a subset of cis-regulatory elements that are likely to be bound by PE-specific TFs. Interestingly, we have found that most of the chromatin accessibility changes happen outside promoters, in regions losing and acquiring enhancer-specific histone marks; conversely, at the promoter level the prominent effect is a change in the histone mark signatures rather than a change in chromatin openness. By leveraging and combining the high-throughput datasets we generated, we have inferred a differential GRN describing the activation of a PE-specific transcriptional program. Apart from confirming the role of known PE markers and the importance of Retinoic Acid receptors, this GRN also highlighted new TF candidates that are likely to be active players in this transcriptional network. To note, TBX1, appears to be regulated through the binding of several upstream TFs to promoter and enhancer regions. Other TFs that are predicted to be activated by upstream factors and, once expressed, to act on their targets to guide the PE specification include HOXA1, HOXB1 and HOXB2, whose expression is controlled by enhancers bound by RARs and by other factors which are not directly activated by RA. Leveraging on these important findings, follow-up mechanistic experiments will be necessary in order to establish a causal relation and to validate such predicted interactions. Overall, our platform represents a powerful resource that can be used at large by the scientific community to study the molecular function of known factors that, despite being crucial for the PE development, have not yet been characterized in the proper cell context, and whose malfunction in this stage is responsible for the progress of several developmental diseases. In addition to this, it also provides a list of new putative cis-regulatory elements that are likely responsible for the activation of important developmental genes and of genes involved in the formation of pharyngeal organs (e.g., thymus, thyroid, parathyroid glands), which can be further validated and characterized. Lastly, our PE cells represent an important developmental intermediate stage that can be further manipulated to produce and engineer pharyngeal organs *in vitro*, with a paramount clinical relevance for precision medicine.

## METHODS

### Cell culture conditions

H9 hESCs (Thomson et al., 1998) were routinely propagated feeder-free in mTeSR1 on cell culture plates previously coated with Matrigel (Corning, cat. n. 354230) following the manufacturer instructions. Undifferentiated hESCs were propagated and passed at least 3 times after thawing and plated for the differentiation when at 80% of confluence. Cells were maintained in culture and expanded at high quality with particular care to avoid any spontaneous differentiation, which would confound downstream differentiation. The day before the induction of the differentiation, hESCs were washed twice in PBS (Cat. n. Gibco 10010-023), dissociated with Accutase (Innovative Cell Technologies, Cat. n. AT104), plated in mTeSR1 + Rock Inhibitor (Y27632, 5uM) to promote cell survival and incubated for 12 hours at 37 °C, 5% CO_2_ for 12 hours. The day after, 30% confluent hESCs were induced to differentiate into definitive endoderm, by incubation with Medium A (Gibco cat. n. A30621-01) for 24 hours and with Medium B (Gibco cat. n. 30624-01) for the following 24 hours at 37 °C, 5% CO_2_. The third day DE was patterned into AFG by the addition of CDM2 medium with A-83-01, 1uM and DM3189, 250nM. The composition of CDM2 basal medium (Loh et al., 2014) was as follows: 50% IMDM (+GlutaMAX, +HEPES, +Sodium Bicarbonate; Gibco, 31980-097) + 50% F12 (+GlutaMAX; Gibco, 31765-092) + 1 mg/mL polyvinyl alcohol (Sigma, P8136-250G) + 1% v/v concentrated lipids (Gibco, 11905-031) + 450 µM monothioglycerol (Sigma, M6145) + 0.7 µg/mL insulin (Roche, 1376497) + 15 µg/mL transferrin (Roche, 652202) and incubated at 37 °C, 5% CO_2_. From day four to day 7, Retinoic Acid 200 nM was added at the CDM2 medium with A-83-01, 1 uM and DM3189, 250 nM and medium was changed every 24 hours.

### RNA isolation and quantitative-RT PCR (qRT-PCR)

Total RNA from hESCs, DE, AFE, and PE cells was isolated by collecting cells using TRI Reagent (Zymo Research), followed by column purification and DNAse treatment using Direct-zol RNA MiniPrep Kit (Zymo Research), and quantified by Nanodrop (Thermo Scientific). RNA (0.5-1.0μg) was reverse transcribed using PrimeScript Reagent Kit (Takara) according to manufacturer’s instructions for quantitative RT-PCR analyses. Quantification analyses were carried out using PowerUp SYBR-Green MasterMix (Thermo Fisher Scientific).

### Bulk RNA-Seq experiment

RNA library preparations and sequencing reactions were conducted at GENEWIZ, Inc/Azenta US, Inc. (South Plainfield, NJ, USA). 1 ug of RNA from three biological replicates of hESCs, DE, AFE and PE conditions was quantified using Qubit 2.0 Fluorometer (Life Technologies, Carlsbad, CA, USA) and RNA integrity was checked using Agilent TapeStation 4200 (Agilent Technologies, Palo Alto, CA, USA). RNA sequencing libraries were prepared using the NEBNext Ultra RNA Library Prep Kit for Illumina using manufacturer’s instructions (NEB, Ipswich, MA, USA). Briefly, mRNAs were initially enriched with Oligod(T) beads. Enriched mRNAs were fragmented for 15 minutes at 94 °C. First strand and second strand cDNA were subsequently synthesized. cDNA fragments were end-repaired and adenylated at 3’ ends, and universal adapters were ligated to cDNA fragments, followed by index addition and library enrichment by PCR with limited cycles. The sequencing library was validated on the Agilent TapeStation (Agilent Technologies, Palo Alto, CA, USA), and quantified by using Qubit 2.0 Fluorometer (Invitrogen, Carlsbad, CA) as well as by quantitative PCR (KAPA Biosystems, Wilmington, MA, USA). The sequencing libraries were clustered on a single lane of a flow cell. After clustering, the flow cell was loaded on an Illumina HiSeq 4000 instrument according to the manufacturer’s instructions. The samples were sequenced using a 2×150 bp paired-end (PE) configuration. Image analysis and base calling were conducted by the HiSeq Control Software (HCS). Raw sequence data (.bcl files) generated from Illumina HiSeq was converted into fastq files and de-multiplexed using Illumina’s bcl2fastq v2.17 software. One mismatch was allowed for index sequence identification. On average, 33 million read pairs were obtained for each sample.

### Bulk RNA-Seq data analysis

Adapter sequences and poor quality ends were removed using the Trimmomatic v0.39 software (Bolger et. al., 2014) with parameters *ILLUMINACLIP:/path/to/adapter:2:30:10:1:true SLIDINGWINDOW:20:15 MINLEN:36*. To produce coverage tracks, reads first were mapped to human GRCh38 genome and GENCODE v35 transcriptome (Frankish et al., 2019) using STAR v2.7.6a software (Dobin et al., 2013), with parameters *--peOverlapNbasesMin 10 --outSAMstrandField intronMotif --outFilterIntronMotifs RemoveNoncanonical --outSAMattrIHstart 0 --outSAMtype BAM SortedByCoordinate*; SAMtools v1.11 merge tool (Li et al., 2009) was used to pool together read alignments from biological replicates, thus producing a single pooled BAM file for each condition; deepTools v3.5.1 (Ramírez et al., 2014) bamCoverage tool was then employed to convert pooled BAM files to bigWig files while removing reads mapping to ENCODE Blacklist regions (Amemiya et al., 2019), with RPKM normalization and the genomic bin size set to 10 bp. Salmon v1.3.0 tool (Patro et al., 2017) was employed to quantify transcript expression, producing isoform-level transcripts per million (TPM) values from a full decoy transcriptome index created using the GENCODE v35 transcriptome and the hg38 genome. Tximport v1.18.0 R package (Soneson et al., 2015) was employed to obtain gene-level TPMs and estimated counts. Such counts were used for the differential gene expression analysis, performed using the DESeq2 v1.30.0 R package (Love et al., 2014), after removing genes with TPM < 1 in at least 10 samples. For each contrast, differential gene expression analysis with independent filtering was run twice, by setting the *lfcThreshold* for the Wald test either to 0 or to 0.58, thus producing a relaxed and a strict set of differentially expressed genes (the FDR threshold was set to 0.01 in both cases); only genes with an average TPM > 1 in at least one of the two conditions under comparison were retained. Apeglm (Zhu et al., 2019) method was employed for log2(FC) shrinkage. Regularized-log (rlog) transformation was applied to count data for subsequent clustering analysis and to produce the heatmap showing the expression of the DEGs identified *via* the strict test. The principal component analysis (PCA) plot was drawn using the DESeq2 “plotPCA” function; the sample-to-sample euclidean distance heatmap was produced using the pheatmap v1.0.12 R package (available at https://CRAN.R-project.org/package=pheatmap). The UpSet plot showing the intersections between DEGs identified in each contrast was drawn using the UpSetR v1.4.0 R package (Conway et al., 2017). The gene expression heatmaps were generated using the ComplexHeatmap v2.6.2 (Gu et al., 2016) R package.

### Comparison of Bulk RNA-Seq samples with mouse scRNA-Seq samples

The count matrix relative to the scRNA-Seq data from embryonic mouse foregut endoderm produced by Han and colleagues was downloaded from the Gene Expression Omnibus GEO archive (GSE136689) (Edgar et al., 2002). Counts were transformed to counts per million values (CPM), and average CPM values were calculated for each gene in each endormal cell cluster. For each cluster, we retrieved the top 10 transcription factor markers from the original publication. The similarity of each cluster with our bulk RNA-Seq samples was assessed by computing the Spearman correlation coefficient between the log10-transformed CPM values of its top TF markers and the log10-transformed TPM values of their human counterparts. Mouse-Human orthology relationships were retrieved from Ensembl 101 database (Howe et al., 2021). The Spearman correlation matrices and the HOX gene expression heatmaps were plotted using the ComplexHeatmap R package.

### scRNA-Seq experiment

For single cell RNA-Seq experiments, DE and PE differentiated cells were washed twice with CDM2 medium to remove dead cells and detached using Accutase. Cells were collected and counted using Countess® II Automated Cell Counter. 500,000 cells for each condition were collected in a PBS-0.04%BSA buffer and processed according to the 10X Genomics Single Cell Protocols Cell Preparation Guide (available at https://assets.ctfassets.net/an68im79xiti/56DlUZEsVOWc8sSG42KQis/35cbcf6dcd4b0c0196263ee93815b0ae/CG000053_CellPrepGuide_RevC.pdf). For each cell type, 5000 cells were loaded per lane on the 10x Genomics Chromium platform, with the goal of capturing 2500 cells. Cells were then processed for cDNA synthesis and library preparation using 10X Genomics Chromium Version 2 chemistry (catalog number 120234) as per the manufacturer’s protocol. cDNA libraries were checked for quality on the Agilent 4200 Tape Station platform and their concentration was quantified by KAPA qPCR. Libraries were sequenced using an Illumina HiSeq 4000 instrument to a depth of, at a minimum, 70,000 reads per cell. For hESCs, sequencing data were previously produced from our lab and are available in the GEO repository with the accession number GSE157475.

### ScRNA-Seq data analysis

Illumina base call files were converted to FASTQ files using the Cell Ranger v2.0 program (Zheng et al., 2017). FASTQ files were then aligned to the hg19 human reference genome using Cell Ranger. The Scanpy v1.7.2 Python package (Wolf et al., 2018) was used for subsequent analyses. Cells from all the three samples were first combined into a single “anndata” object, containing 6115 hESCs cells, 2566 DE cells and 2525 PE cells. Quality control metrics, including the number of detected genes per cell, the total counts per cell and the percentage of counts belonging to mitochondrial genes were calculated using the function “scanpy.pp.calculate_qc_metrics”. We first filtered out low-quality cells that expressed fewer than 13,000, 20,000 and 15,000 counts for hESCs, DE and PE, respectively. These thresholds were chosen based on the distribution of the total counts for each sample. We also excluded cells that expressed more than 7,000 genes (which would imply doublets) or that expressed more than 10% mitochondrial genes (indicative of dead cells in this dataset) (Luecken & Theis, 2019). Finally, we filtered out cells with less than 2500 expressed genes using the function “scanpy.pp.filter_cells” and genes expressed in less than 10 cells using the function “scanpy.pp.filter_genes”. After quality control, we obtained 4887 hESCs cells, 1127 DE cells and 1115 PE cells. Next, normalization was performed by dividing raw counts by the library counts sum and multiplying by a factor of 100,000, using the function “scanpy.pp.normalize_total” with *target_sum*parameter set to 100000. After log normalization, the highly variable genes were selected with the function “scanpy.pp.highly_variable_genes”, with *max_mean*, *min_mean* and *min_disp* parameters set to 5, 0.0125 and 0.5, respectively, obtaining 2215 genes. From this set we removed ribosomal genes, finally obtaining 2203 highly variable genes. The total counts and the percentage of mitochondrial counts were regressed out as potential confounding factors with the function “scanpy.pp.regress_out”. Genes were then scaled to zero mean and unit variance, clipping maximum values to 10 (“max_value” parameter). A principal component analysis was performed on the scaled matrix with the function “scanpy.tl.pca”, using the “arpack” singular value decomposition solver, and a k-nearest neighbor graph was computed with the function “scanpy.pp.neighbors”. Dimensionality reduction through the Uniform manifold approximation and projection (UMAP) (Becht et al., 2019) algorithm was performed with the function “scanpy.tl.umap” using the highly-variable genes and initializing the positions using the Partition-based graph abstraction (PAGA) (Wolf et al., 2018). Cell clustering was performed with the Leiden algorithm using the function “scanpy.tl.leiden” with default parameters, obtaining 10 clusters: 7 clusters of hESCs, 2 of DE and 1 of PE cells.

Differentially expressed genes between clusters were identified with the function “scanpy.tl.rank_genes_groups” using the t-test; for each group, the top 50 DEGs were chosen based on the Z-score returned by this function. All single-cell RNA-Seq plots were also generated using Scanpy.

### Retrieval of transcription start sites and promoter regions

TSSs of protein-coding transcripts with annotated 5’UTR were retrieved from Refseq v109.20211119 curated annotation (O’Leary et al., 2016) and from the “upstream1000.fa” file provided by the UCSC Genome Browser (Kent et al., 2002) (available at https://hgdownload.soe.ucsc.edu/goldenPath/hg38/bigZips/). The hg38 genomic coordinates of such TSSs were extended by 3000 bp both upstream and downstream to obtain a set of promoter regions. These promoters were assigned to their corresponding GENCODE protein-coding genes using the BEDTools intersect tool. Promoters of protein-coding genes that do not produce polyadenylated transcripts were not kept for further analyses; such genes were defined as non-expressed genes (TPM < 1 in at least 10 Bulk RNA-Seq samples) that do not overlap with any poly(A) feature from PolyASite 2.0 database (Herrmann et al., 2020) and GENCODE annotation (available at https://www.gencodegenes.org/human/release_35.html).

### ATAC-Seq experiment

ATAC-Seq library preparation and sequencing reactions were conducted at GENEWIZ, Inc/Azenta US, Inc. (South Plainfield, NJ, USA). DE, AFE and PE live cell samples (two biological replicates per cell type) were thawed, washed, and treated with DNAse I (Life Tech, Cat. #EN0521) to remove genomic DNA contamination. Live cell samples were quantified and assessed for viability using a Countess Automated Cell Counter (ThermoFisher Scientific, Waltham, MA, USA). After cell lysis and cytosol removal, nuclei were treated with Tn5 enzyme (Illumina, Cat. #20034197) for 30 minutes at 37°C and purified with Minelute PCR Purification Kit (Qiagen, Cat. #28004) to produce tagmented DNA samples. Tagmented DNA was barcoded with Nextera Index Kit v2 (Illumina, Cat. #FC-131-2001) and amplified via PCR prior to a SPRI Bead cleanup to yield purified DNA libraries. The sequencing libraries were clustered on one lane of a flow cell. After clustering, the flow cell was loaded on an Illumina HiSeq 4000 instrument according to the manufacturer’s instructions. The samples were sequenced using a 2×150 bp PE configuration. Image analysis and base calling were conducted by the HiSeq Control Software (HCS). Raw sequence data (.bcl files) generated from Illumina HiSeq was converted into fastq files and de-multiplexed using Illumina’s bcl2fastq v2.20 software. One mismatch was allowed for index sequence identification. On average, ∼98 million read pairs were obtained for each sample.

### ATAC-Seq data analysis

Sequencing adapters and low-quality bases were trimmed using the Trimmomatic v0.39 software with parameters *ILLUMINACLIP:/path/to/adapter:2:30:10:1:true SLIDINGWINDOW:20:15 MINLEN:36*. Preprocessed reads were then aligned to the hg38 genome using Bowtie2 (Langmead and Salzberg, 2012) with parameters *--wrapper basic-0 --fr -X 2000*. Aligned reads were filtered using SAMtools to keep only concordant primary alignments having a minimum mapping quality of 30. PCR or optical duplicates were marked using Picard v2.25.1 tool (available at https://broadinstitute.github.io/picard/) and removed. Reads mapping to mitochondrial DNA and to unplaced contigs were filtered out. Aligned reads were also shifted as in (Buenrostro et al., 2013) using the deepTools alignmentSieve tool with the *--ATACshift* parameter. After this shift, reads falling in ENCODE Blacklist regions were removed using BEDTools v2.30.0 pairToBed tool (Quinlan & Hall, 2010). Read alignments from biological replicates were pooled together using SAMtools merge. deepTools bamCoverage tool was then employed to convert BAM files (both from individual and pooled replicates) to bigWig files with RPGC normalization and the genomic bin size set to 10 bp (for track visualization) and to 50 (for coverage heatmaps).

MACS2 v2.2.7.1 callpeak tool (Zhang et al., 2008) was used to identify open chromatin regions in each replicate, with parameters *-f BAMPE --call-summits -g hs --keep-dup all*. Peaks identified by MACS2 in all the samples were used to determine a consensus peak set using the “dba” function from the DiffBind v3.0.13 R package (Ross-Innes et al., 2012), setting the *minOverlap* parameter to 2. Reads mapping in 201 bp intervals centered on consensus peak summits were counted using the “dba.count” function, with the *filter* parameter set to 0; counts were normalized using full library size with the “dba.normalize” function. PCA was drawn using the “dba.plotPCA” function; Pearson correlation coefficient values, calculated on the normalized read counts between each pair of samples using the “dba.plotHeatmap” function, were employed to draw a sample-to-sample distance matrix using the pheatmap R package. Only consensus peaks called by MACS2 in both replicates of at least one condition were employed to draw the Venn diagram, produced using the BioVenn v1.1.3 R package (Hulsen et al., 2008). Differential accessibility analysis was performed for each contrast with the “dba.analyze” function, setting the underlying method to DESeq2; a paired design, justified by the timing of sample preparation and sequencing, was employed only for the DE vs PE contrast. For each A vs B contrast, we identified three classes of 201 bp peaks:

- Common peaks, called by MACS2 in both replicates of A and/or B and with DiffBind FDR > 0.01 and/or absolute log2(FC) < 1;
- Lose peaks, called by MACS2 in both replicates of A and with DiffBind FDR < 0.01 and log2(FC) < −1;
- Gain peaks, called by MACS2 in both replicates of B and with DiffBind FDR < 0.01 and log2(FC) > 1.

The heatmap showing the clustering of differentially accessible regions (DARs) was produced using the ComplexHeatmap R package. PhyloP basewise conservation scores derived from Multiz alignment of 100 vertebrate species (Pollard et al., 2010) were retrieved for DARs and Common peaks using the GenomicScores v2.2.0 R package (Puigdevall & Castelo, 2018).

Heatmaps and profile plots of ATAC-Seq signal around the TSSs of protein-coding genes were drawn by applying the deepTools computeMatrix, plotHeatmap and plotProfile tools to the previously produced BigWig files with 50 bp resolution.

The protein-coding promoter chromatin accessibility status was evaluated by searching for overlaps between protein-coding promoters and consensus ATAC-Seq peaks using BEDTools intersect. Non-promoter peaks were identified based on the absence of overlap with any TSS +- 3kb region derived from RefSeq and UCSC knownGene annotation. The proximity of any consensus ATAC-Seq peak to protein-coding gene TSSs was evaluated using the BEDTools closest tool.

### Transcription factor motif and footprint analyses

The TF motifs used in the present work are those composing the non-redundant, clustered gimme.vertebrate.v5.0 database, which is available within the GimmeMotifs v0.17.0 analysis framework (Bruse & van Heeringen, 2018). To identify differentially enriched motifs among DE, AFE and PE, we first collected all DARs and calculated the average of the normalized counts across the biological replicates for each condition; such mean accessibility measurements were then log2-transformed and subsequently centered by subtracting the mean of the log2-transformed values across the three conditions. The resulting table of scaled read counts was provided to the GimmeMotifs maelstrom tool, which was run with the *--no-filter* option. This tool combines different motif enrichment methods to calculate, for each TF motif, a set of condition-specific combined Z-scores, each one representing the enrichment of the motif among the condition-specific accessible regions.

TF footprints (FP) were individually identified for DE, AFE and PE conditions by applying the 2017-04-27 version of the PIQ tool (Sherwood et al., 2014) to pooled BAM files, using the “pairedbam2rdata.r” script to convert them to internal binary format, setting the purity score threshold to 0.7 and using the gimme.vertebrate.v5.0 motif file, after converting it to JASPAR format (Sandelin et al., 2004) with UniversalMotif v1.8.3 R package (available at https://bioconductor.org/packages/universalmotif/), as input motif database for the “pwmmatch.r” script; only motifs belonging to TF with average TPM > 5 in at least condition were used in this analysis.

FP enrichment analysis was performed using the BiFET tool. Specifically, for each A vs B differential accessibility contrast, BiFET was employed to evaluate the enrichment of FPs identified in B among the Gain peaks and the enrichment of FPs identified in A among the Lose peaks, using the Common peaks as background loci in both cases. The normalized read counts and the GC content of each ATAC-Seq consensus peak, which are employed by BiFET for bias correction, were calculated using DiffBind and HOMER tools (Heinz et al., 2010), respectively. “findOverlaps” function from the GenomicRanges v1.42.0 R package (Lawrence et al., 2013) was used to find the FPs overlapping consensus peaks.

For the integrated analysis of TF activity, only motifs with an absolute maelstrom Z-score >= 2 in at least one condition, a BiFET adjusted p-value < 0.001 in at least one set of DARs and an average transcription factor gene TPM > 5 in at least one cell type were initially selected. Furthermore, only TF genes found to be differentially expressed in at least one contrast were employed. BiFET adjusted p-values were converted to −log10(adjusted p-values), after replacing 0 values with 1×10^-16^ to avoid infinite numbers; the average of these transformed p-values was computed for each cell-type specific set of DARs, thus obtaining a FP enrichment score for each condition. Z-scores of log2-transformed average TPMs along the cell types were also computed to obtain a set of condition-specific expression Z-scores for each TF. The final set of cell type-specific TF motifs was obtained by selecting only motifs whose maelstrom Z-scores are positively correlated (Pearson correlation coefficient > 0.5) with the FP enrichment scores and with the expression Z-scores. The heatmap showing the enrichment of these motifs was drawn using the ComplexHeatmap R package.

### ChIP-Seq experiment

ChIP experiments were performed on chromatin extracts according to the manufacturer’s protocol (MAGnify ChIP, Life Technologies Cat. n. 492024). For each immunoprecipitation reaction, 10ug of sheared chromatin from DE and PE differentiated cells was used (two biological replicates per ChIP experiment). Sheared chromatin was incubated O.N. with 5μg of anti-H3K27me_3_ (Abcam Cat. n. ab6002), H3K4me_3_ (Active Motif Cat. n. 39159), H3K27ac (Active Motif Cat. n. 39133), or H3K4me_1_ (Abcam ab8895) antibodies. ChIP-Seq library preparation and sequencing reactions were conducted at GENEWIZ, Inc/Azenta US, Inc. (South Plainfield, NJ, USA). Immunoprecipitated (IP) and input DNA samples were quantified by Qubit 2.0 Fluorometer (Invitrogen, Carlsbad, CA) and the DNA integrity was checked with 4200 TapeStation (Agilent Technologies, Palo Alto, CA, USA). NEBNext Ultra DNA Library Preparation kit was used following the manufacturer’s recommendations (Illumina, San Diego, CA, USA). Briefly, the ChIP DNA was end-repaired and adapters were ligated after adenylation of the 3’ ends. Adapter-ligated DNA was size selected, followed by clean up, and limited cycle PCR enrichment. The ChIP library was validated using Agilent TapeStation and quantified using Qubit 2.0 Fluorometer as well as real time PCR (Applied Biosystems, Carlsbad, CA, USA). The sequencing libraries were multiplexed and clustered on one lane of a flow cell. After clustering, the flow cell was loaded on an Illumina HiSeq 4000 instrument according to the manufacturer’s instructions (Illumina, San Diego, CA, USA). Sequencing was performed using a 2×150 bp PE configuration. Image analysis and base calling were conducted by the HiSeq Control Software (HCS). Raw sequence data (.bcl files) generated from Illumina HiSeq was converted into fastq files and de-multiplexed using Illumina’s bcl2fastq v2.17 software. One mismatch was allowed for index sequence identification. On average, 49 million read pairs were obtained for each sample.

### ChIP-Seq data analysis

The preprocessing, alignment and post-alignment filtering of reads, as well as the generation of bigWig files with RPGC normalization, were performed as in the ATAC-Seq data analysis, except for the read alignments shift, which was skipped. 50 bp resolution bigWig files for individual replicates of immunoprecipitated samples were given as input to deepTools multiBigwigSummary to compute average RPGC scores for 10 kb genomic bins; deepTools plotCorrelation was employed on the resulting output file to produce the Pearson correlation matrix and the hierarchical clustering of samples. In addition, we compared the 50 bp resolution pooled coverage tracks of IP and input samples using deepTools bigwigCompare to generate BigWig files reporting the log2(FC) of the IP signal over the input for each 50 bp genomic bin. deepTools computeMatrix, plotHeatmap and plotProfile tools were applied to these files to draw heatmaps and profile plots of ChIP/input signal around the TSSs of protein-coding genes and the summits of ATAC-Seq consensus peaks. For the former analysis, TSSs were stratified based on the overlap with ATAC-Seq peaks, evaluated using BEDTools intersect after replacing the boundaries of the DiffBind consensus peaks with those of the corresponding merged MACS2 peaks.

Chromatin state discovery was performed using the ChromHMM v1.22 software (Ernst & Kellis, 2012). As a first step, all the BAM files were binarized with the BinarizedBam module, using a bin size of 200 bp. Concatenated model learning was conducted with the LearnModel module, using the input samples as control data to adjust the binarization threshold locally. This module was employed to build models with a number of states ranging from 6 to 16. We decided to focus on a model with 10 states for the subsequent analyses, since it delivered a compact and meaningful representation of the main chromatin states that can be produced with the 4 histone marks under analysis. The LearnModel module produced 200 bp chromatin state calls for DE and PE cell types. OverlapEnrichment module was employed to compute, both for DE and PE chromatin state annotations, the fold enrichment relative to a set of genomic features derived from the RefSeq annotation and to the ATAC-Seq consensus peaks, classified based on the DE vs PE contrast. The fold enrichment of chromatin states relative to the neighborhood of TSSs derived from RefSeq annotation was computed with the NeighborhoodEnrichment module. The relationship between accessible regions and chromatin states was investigated by using BEDTools intersect to assign ATAC-Seq consensus peaks to the 200 bp genomic bins containing their summits. Protein-coding TSSs were assigned to their corresponding bins using the “findOverlaps” function from the GenomicRanges R package.

The state transition enrichment analysis was inspired by a work by Fizier and colleagues (Fiziev et al., 2017). Specifically, we first calculated the number of 200 bp bins involved in each possible chromatin state transition from DE to PE; for each transition, we also calculated the expected number of transitioning bins as the average of the number of transitions obtained after shuffling the state calls 1000 times; we then divided the observed counts by the expected counts to compute an enrichment score for each transition, thus controlling for the state coverage; finally, fold enrichment values were obtained by dividing the enrichment score of each transition by the enrichment score of the transition having opposite direction, thus controlling for the overall similarity between the two states involved in the transition.

For each genomic bin, the nearest expressed protein-coding gene (average TPM > 1 in DE and/or in PE) with a TSS within a distance of 50 kb, if any, was identified using the “distanceToNearest” function from the GenomicRanges R package. To test the association between chromatin state transitions and deregulation of nearby genes, for each state transition we calculated the number upregulated, downregulated and non-differentially expressed genes in the proximity of the bins undergoing the transition (*UP_T_*, *DOWN_T_*, *NO_T_*) and of all the other bins (*UP_O_*, *DOWN_O_*, *NO_O_*), used as controls (also transitions between identical states were employed in this analysis). Then, three Fisher’s exact tests were performed for each transition (the numbers within the square brackets representing a row of a 2×2 contingency table), obtaining a set of p-values (adjusted using the Benjamini–Hochberg procedure):

- *P_UPDOWN_*: [*UP_T_*, *DOWN_T_*] vs [*UP_O_*, *DOWN_O_*];
- *P_UP_*: [*UP_T_*, (*DOWN_T_*+*NO_T_*)] vs [*UP_O_*, (*DOWN_O_*+*NO_O_*)];
- *P_DOWN_*: [*DOWN_T_*, (*UP_T_*+*NO_T_*)] vs [*DOWN_O_*, (*UP_O_*+*NO_O_*)].

For each transition, we also computed:

- 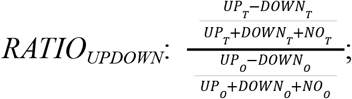
- 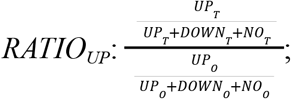
- 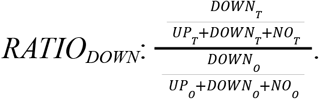

A state transition was considered as enriched in nearby upregulated genes when *RATIO_UPDOWN_* > 1, *RATIO_UP_*> 1.2, *P_UPDOWN_* < 0.05 and *P_UP_* < 0.05 or enriched in nearby downregulated genes when *RATIO_UPDOWN_* < 1, *RATIO_DOWN_* > 1.2, *P_UPDOWN_* < 0.05 and *P_DOWN_* < 0.05.

The association between chromatin state transitions and differential chromatin accessibility was evaluated following a similar procedure, in which the number of upregulated genes was replaced by the number of bins with Gain peaks, the number of downregulated genes was replaced by the number of bins with Lose peaks and the number of non-differentially expressed genes was replaced by the number of genomic bins with no Gain or Lose peaks.

For the FP enrichment analysis of chromatin state-specific DARs, performed using BiFET, we focused on transitions enriched either in Lose or Gain peaks. Lose and Gain peaks were divided based on their chromatin state in DE and PE, respectively. For each class of Lose peaks, we evaluated the FP enrichment of downregulated TFs with an average DE TPM > 5. For each class of Gain peaks, we evaluated the FP enrichment of upregulated TFs with an average PE TPM > 5. In both cases, for each state-specific class of DARs, we used all the Common peaks with a corresponding chromatin state in DE and/or in PE as background regions.

The heatmaps showing the results of the enrichment analyses performed on chromatin state transitions and on state-specific DARs were drawn using the ComplexHeatmap R package.

### Functional enrichment and feature distribution analyses

All the GO Biological Process term enrichment analyses were performed using the Over-Representation Analysis (ORA) method available within the WebGestaltR v0.4.4 R package (Liao et al., 2019). For each analysis we used a different reference set:

- ORA of DEG clusters: all the genes with average TPM > 1 in at least one of the DE, AFE and PE conditions;
- ORA of upregulated protein-coding genes with a non-promoter Gain peak harboring a RARE FP within 50 kb (AFE vs PE contrast): all the expressed protein-coding genes (average TPM > 1 in AFE and/or PE) with an ATAC-Seq consensus peak within 50 kb;
- ORA of protein-coding genes with a TSS transitioning from a bivalent state in DE (TssBiv or EnhBiv) to an active promoter state in PE (TssA, Tss or TssFlnk): all the protein-coding genes with average TPM > 1 in DE and/or PE.

WebGestaltR was employed to perform Gene Set Enrichment Analysis (GSEA), using shrunken log2(FC) values as a ranking metric.

GREAT analysis of each DAR cluster was performed employing the rGREAT v1.22.0 R package (available at https://github.com/jokergoo/rGREAT), using all the DARs and Common peaks as background regions.

The bar plots showing the genomic annotation of of ATAC-Seq peaks were produced using the ChIPseeker v1.26.0 R package (Yu et al., 2015), employing as a TxDb object the one provided by the TxDb.Hsapiens.UCSC.hg38.knownGene v3.10.0 R package (Bioconductor Core Team & Bioconductor Package Maintainer, 2019).

### Gene regulatory network inference

ANANSE v0.3.0+3.g18995f0 software was used to infer GRNs for DE and PE stages and to identify the key TFs in the PE specification process. First, the binding module was employed to predict transcription factor binding individually for each cell type, by providing it with the filtered BAM files obtained from ATAC-Seq and H3K27ac HM CHIP-Seq data, and using the ANANSE REMAP model v1.0 (available at https://zenodo.org/record/4768075/files/ANANSE.REMAP.model.v1.0.tgz), which includes average ChIP-seq signal obtained from the ReMap database (Chèneby et al., 2018); the default motif database (gimme.vertebrate.v5.0) was used in this step. The resulting output files were separately given as input to the network module, thus producing two GRNs, one for DE and one for PE. Finally, the influence module was used to calculate a differential GRN and to compute the influence scores for the transition from DE to PE, providing it with the DE and PE GRNs as the source and target networks, respectively, and with the results of the relaxed differential gene expression analysis performed on the DE vs PE contrast. To increase the number of possible TF-target interactions, which were subsequently filtered during the generation of the PE-specific TF-TF interaction network, the differential GRN was obtained using the top 1,000,000 edges of both source and target networks. To build the PE-specific TF-TF interaction network, the differential GRN was initially filtered to retain only the TF-TF interactions with a differential score > 0.7. To focus on factors specifically active in PE, we only kept interaction involving TFs having average TPM in PE > 5 and which met at least one of the following conditions:

- From the integrated TF activity analysis, the TF was found to be specifically active in the PE stage (all the TFs below TEAD2 in Figure 4A);
- From the FP enrichment analysis of chromatin state-specific DARs, the TF FPs were found to be enriched in at least one class of Gain peaks (BiFET p-value < 0.001);
- The TF was among the 30 top TFs based on ANANSE sumScaled influence score.

Finally, we only kept the interactions in which the source TF had a FP inside a Gain peak (DE vs PE contrast) located at less than 50 kb from the TSS of the target TF. The resulting network was imported in the Cytoscape v3.9.1 software (Shannon et al., 2003), which was used to arrange and visualize it, after filtering out the interactions mediated by Gain peaks located at less than 25 kb from the target to better view the most relevant interactions.

## DATA AVAILABILITY

Bulk RNA-seq, ATAC-Seq and CHIP-seq data associated with this study are available in the GEO repository with the accession numbers GSE206567, GSE206568 and GSE206566, respectively.

## ACKNOWLEDGMENTS

The authors are grateful to the Stanford 10X genomic facility for the scRNAseq experiment and to all the lab members of the Sebastiano and Tartaglia Labs and the Stanford 22q11DS consortium for the scientific discussion.

## AUTHOR CONTRIBUTIONS

V.S. and An.Ci. designed and conceived the study. An.Ci. performed the cellular and molecular experiments with help from D.G. and M.C.. Al.Co. conceived and performed most of the bioinformatics analyses. A.P. performed the RA titration experiments. M.M. performed the single-cell RNA-seq analysis with help from JF. F.B., A.B., and M.G.R. provided help and suggestions in the data interpretation. The paper was written by An.Ci. and Al.Co. with the supervision and major contribution from V.S. and G.G.T., and suggestions from all the other authors.

## SUPPLEMENTARY MATERIAL

Figures S1-S6

