## Supplementary figures for "Multi-omics analysis reveals a crucial role for Retinoic Acid in promoting epigenetic and transcriptional competence of an *in vitro* model of human Pharyngeal Endoderm"

**Supplementary figure 1**

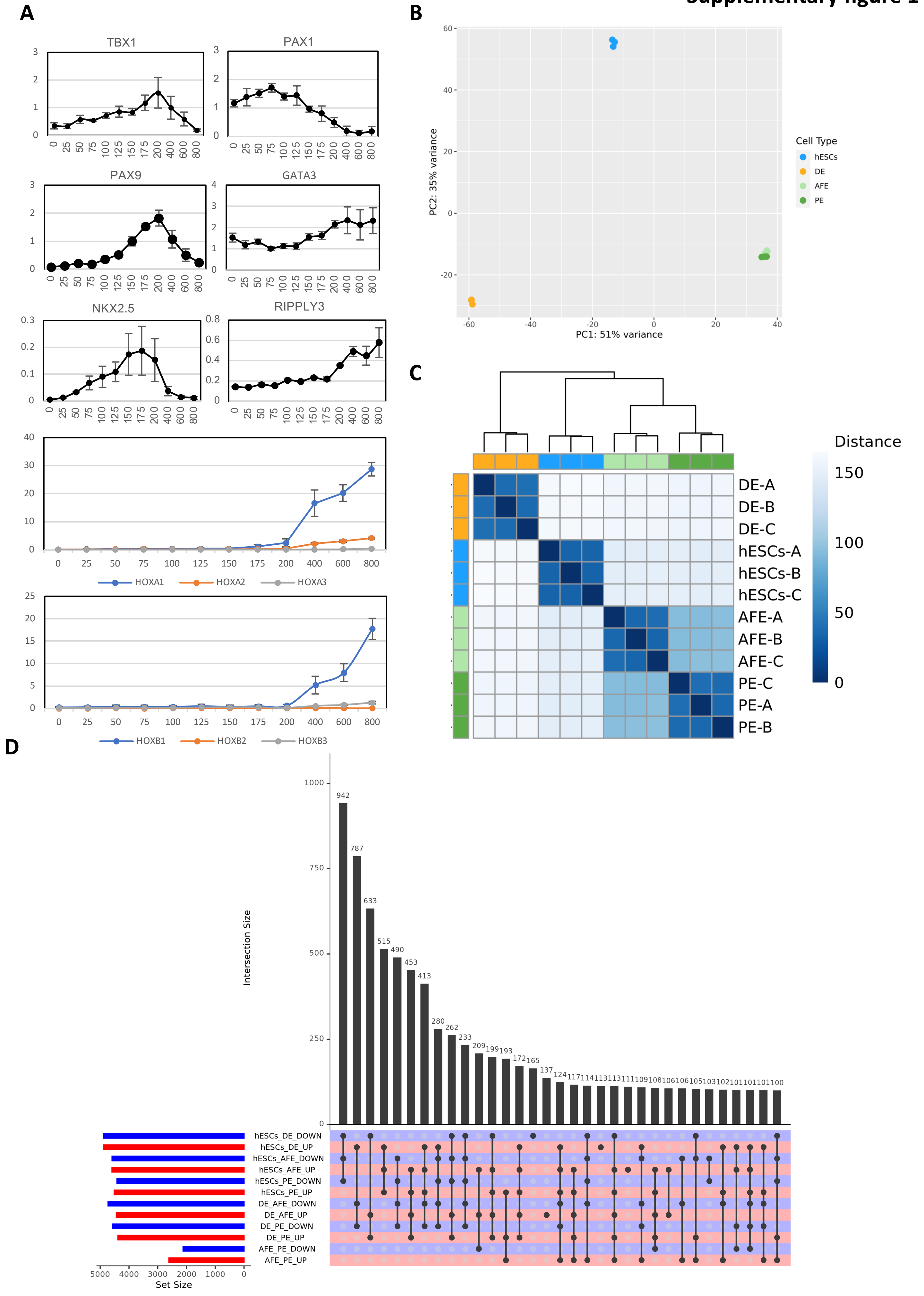

### Supplementary figure 1

#### GO Biological Process

**E**

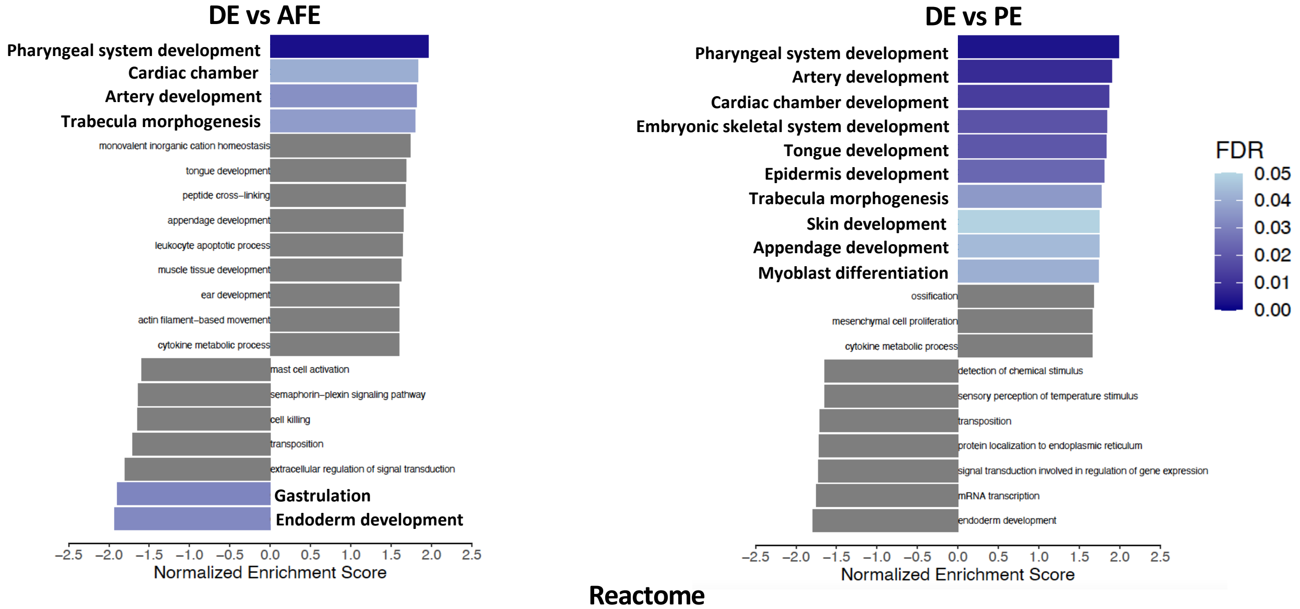


#### Reactome

**F**

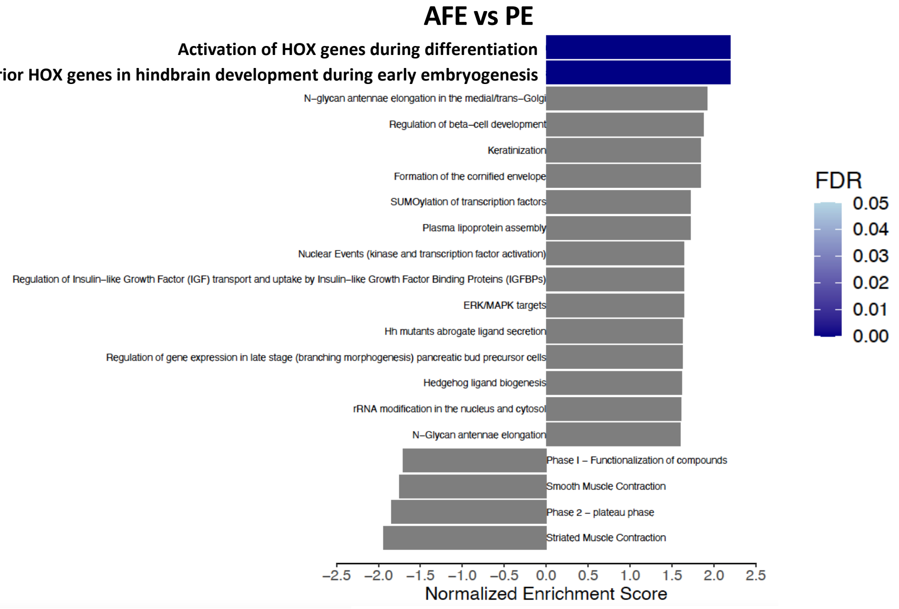


**G**

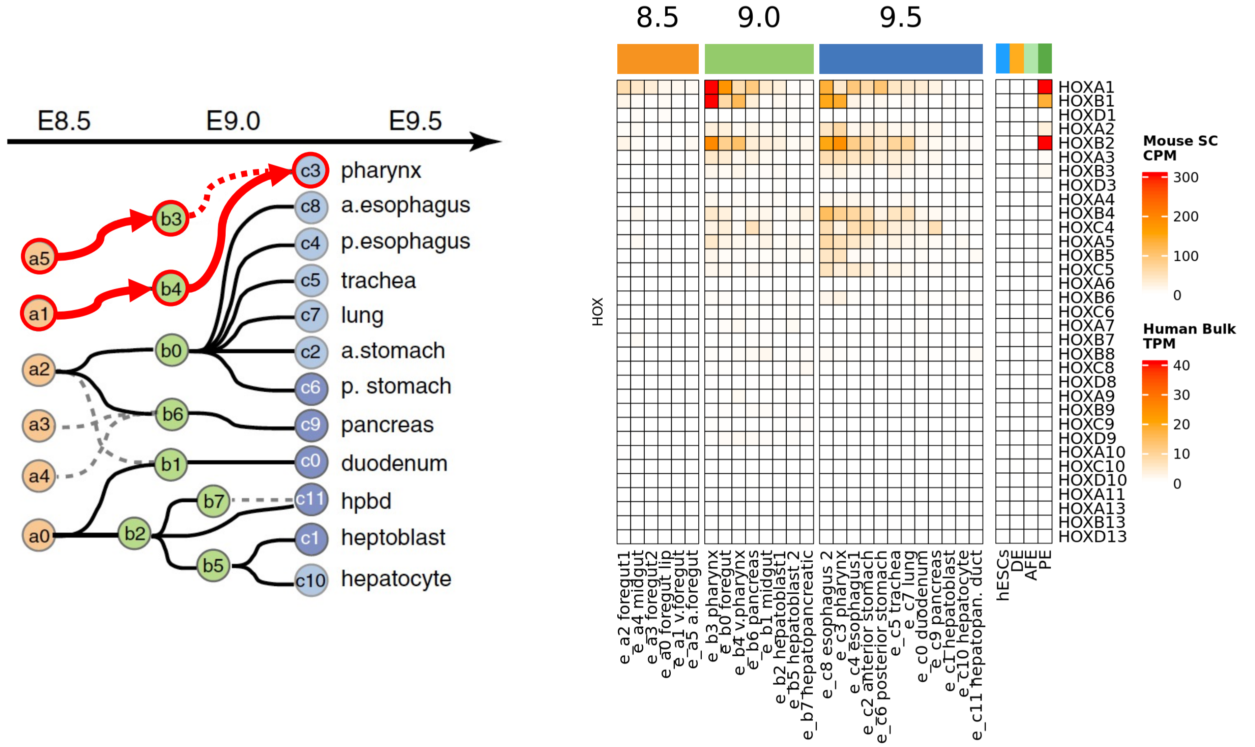

**Supplementary figure 1:**
**(A)** RT-qPCR marker quantification from an *in vitro* differentiation to PE using increasing RA concentration (0-800nM). Data were normalized on PDGB expression and represent means ± SEM of three independent time course experiments.
**(B)** Principal Component Analysis (PCA) plot showing the clustering of hESCs, DE, AFE and PE samples based on their gene expression profile measured via Bulk RNA-Seq of biological triplicates. Principal component 1 (PC1) and 2 (PC2) were identified based on the rlog-tranformed counts relative to the 500 most variable genes.
**(C)** Sample-to-sample euclidean distance heatmap showing the similarity of hESCs, DE, AFE and PE samples based on their gene expression profile measured via Bulk RNA-Seq of biological triplicates. Euclidean distances between rlog-transformed count profiles were also used to draw the dendrogram showing the hierarchical clustering of samples.
**(D)** UpSet plot showing the number of significantly downregulated and upregulated genes (absolute log2[FC] significantly > 0, FDR < 0.01) for all the possible contrasts among hESCs, DE, AFE and PE samples, along with the intersections of different sets consisting of at least 100 genes.
**(E)** Bar plot showing the top 20 functional categories based on absolute Normalized Enrichment Score (NES) calculated performing Gene Set Enrichment Analysis (GSEA) on three contrasts: DE vs AFE and DE vs PE (upper panel), for which enriched non-redundant GO Biological Processes are shown, and AFE vs PE (bottom panel), for which enriched Reactome categories are shown. Positive NESs stand for enrichment among upregulated genes, negative NESs stand for enrichment among downregulated genes. Different shades of blue are used to represent categories with FDR < 0.05, whose names are also highlighted; grey is used to represent categories with FDR >= 0.05.
**(F)** Spatial and temporal developmental tree with lineage prediction of Endodermal Cells during mouse embryonic foregut development at E8.5, E9.0 and E9.5. (adapted from Han et al., 2020). The developmental trajectory of our PE cells based on the analysis shown in figure 1E is highlighted in red.
(**G**) Heatmaps of HOX genes expression profile in the different mouse endodermal cell clusters (data from Han et al., 2020) at different developmental stages (left) and in HESCs, DE, AFE and PE samples (right). Expression levels are shown as gene-level Transcript Per Million (TPM) values.

### Supplementary figure 2

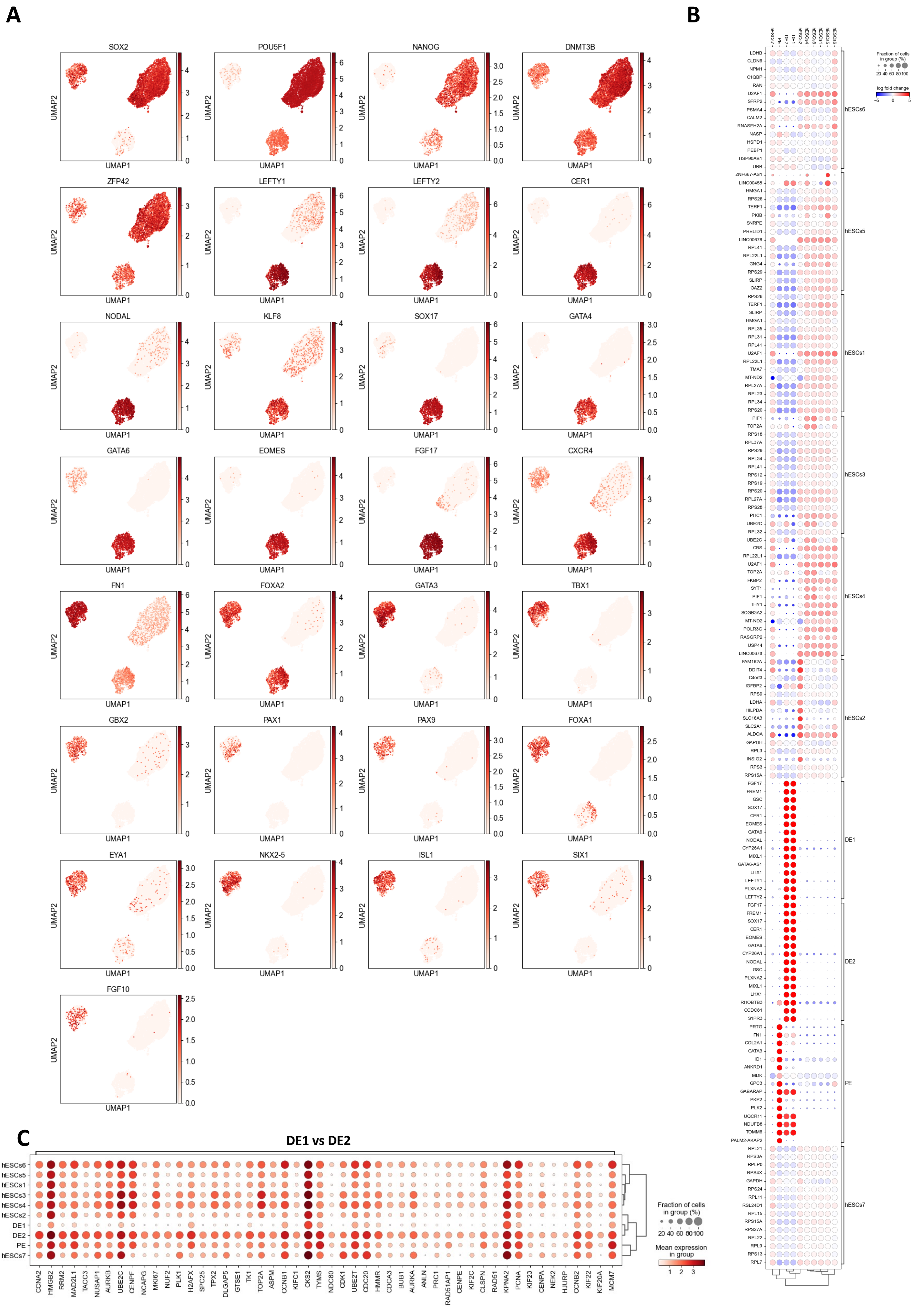

**Supplementary figure 2:**

**(A)** UMAP plot showing the single-cell gene expression (log[norm.counts+1]) of known HESCs, DE, and PE marker genes in HESCs, DE and PE cells.

**(B)** Dot plot showing the top 15 genes responsible for the identity of each cluster and their expression (log[norm.counts+1]) in the 10 different clusters. The dot size represents the percentage of cluster cells expressing the gene and the color represents the log2(FC) of the gene expression compared to the other clusters.

**(C)** Dot plot showing the top 50 DEGs identified between clusters DE1 and DE2 and their expression expression (log[norm.counts+1]) in the 10 different clusters. The dot size represents the percentage of cluster cells expressing the gene and the color represents the average expression in the cluster.

### Supplementary figure 3

A

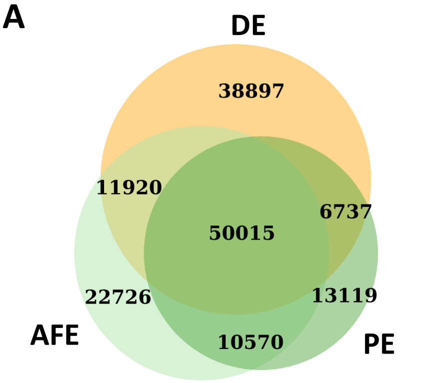


| | Lose | Gain |
|---|---|---|
| DE vs AFE | 41252 | 25553 |
| DE vs PE | 39146 | 19694 |
| AFE vs PE | 20907 | 14870 |

B

C

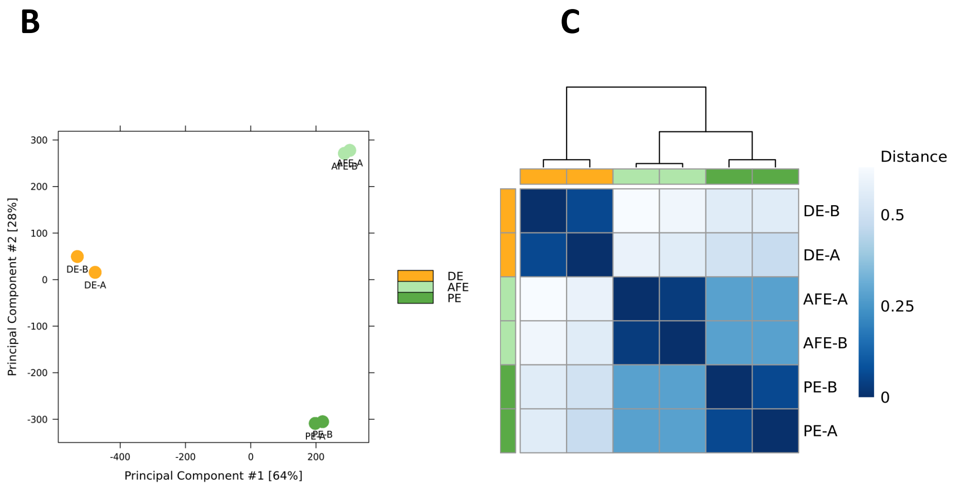


D

E

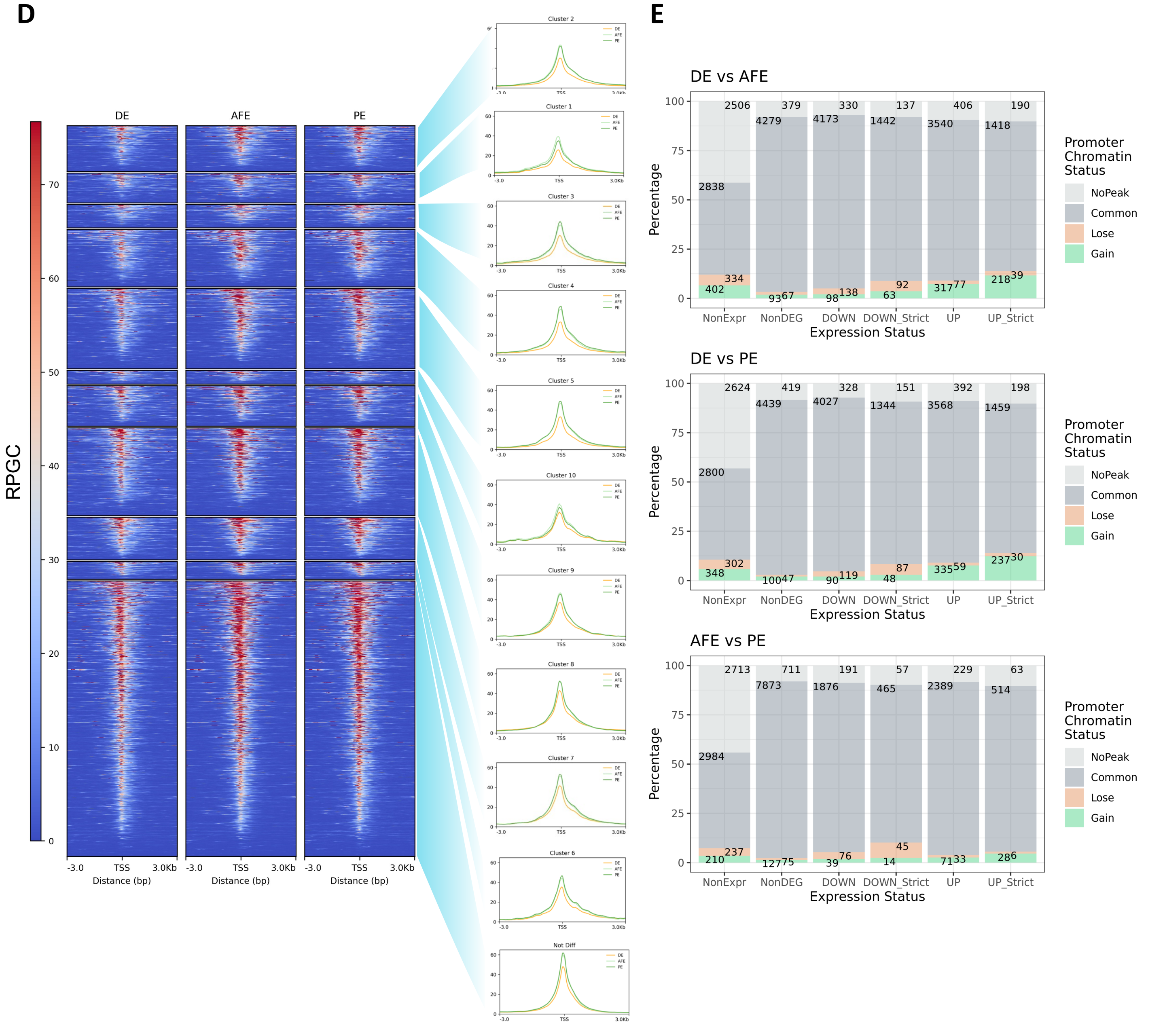

**Supplementary figure 3**

F

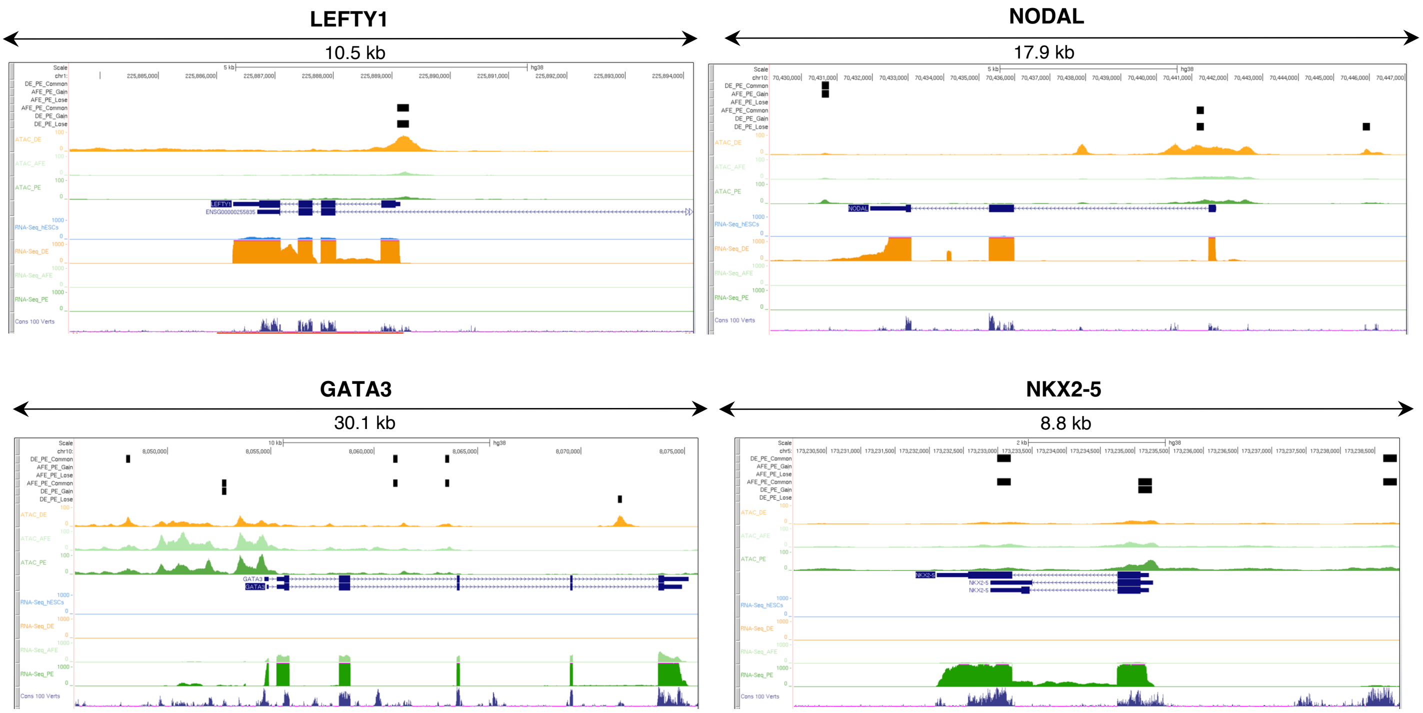


G

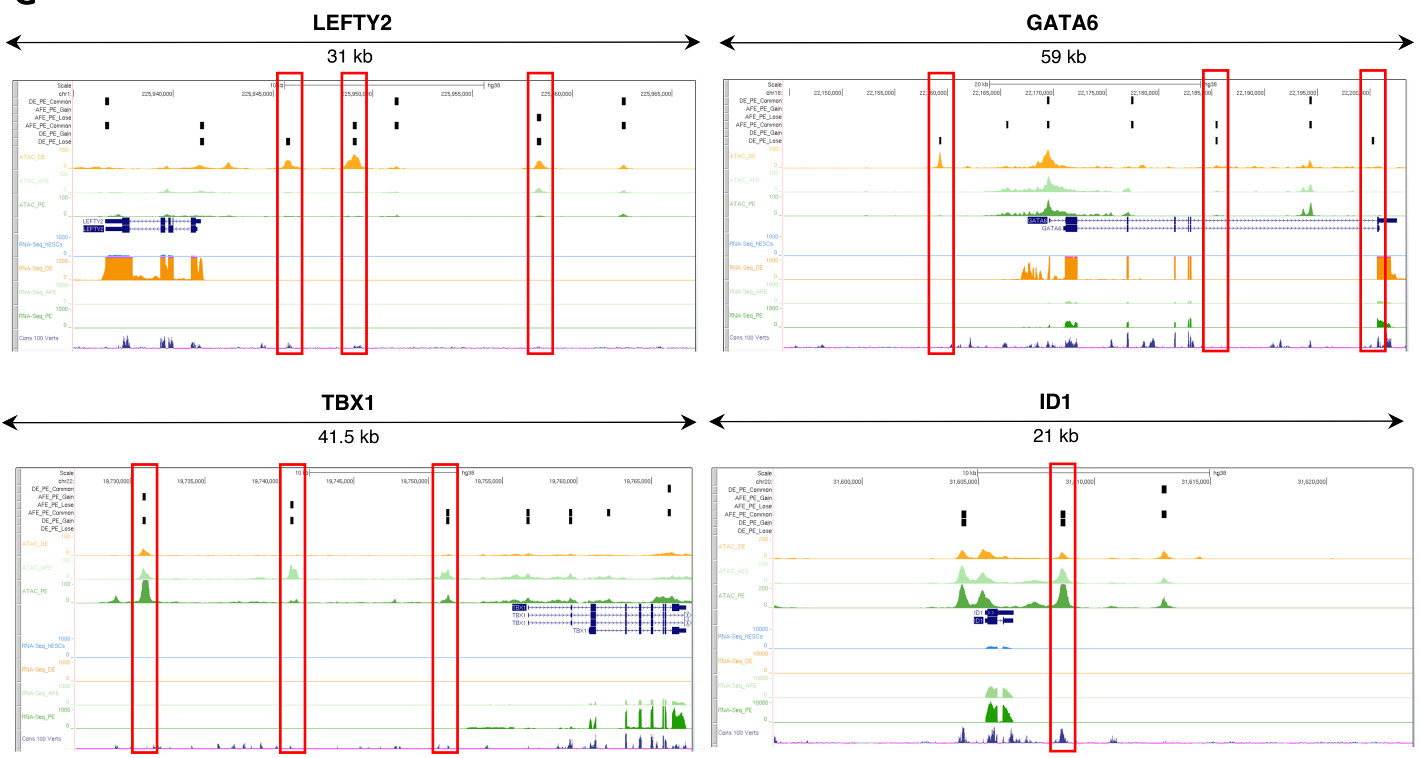

**Supplementary figure 3:**

**(A)** Venn diagram showing the consensus ATAC-Seq peak sets identified in each condition and their overlaps (upper panel). Table showing, for each differential accessibility contrast, the number of Gain (absolute log2[FC] > 1, FDR < 0.01) and Lose (absolute log2[FC] < -1, FDR < 0.01) peaks.

**(B)** PCA plot showing the clustering of DE, AFE and PE samples based on their chromatin accessibility profile measured via ATAC-Seq of biological duplicates. Principal component 1 (PC1) and 2 (PC2) were identified based on the normalized read counts calculated for all peaks.

**(C)** Sample-to-sample Pearson correlation heatmap showing the similarity of DE, AFE and PE based on their chromatin accessibility profile measured *via* ATAC-Seq of biological duplicates. Sample-to-sample distances were calculated as 1-Pearson correlation of the normalized read counts computed for all peaks. Such distances were also used to draw the dendrogram showing the hierarchical clustering of samples.

**(D)** Heatmaps showing DE, AFE and PE ATAC-Seq signal in 6 kb-long regions around the TSSs of protein-coding non-DEGs and DEGs belonging to the clusters shown in Figure 1C. Signal was calculated on merged replicates as Reads Per Genome Coverage (RPGC) values with a bin size of 50 bp. Summary plots reporting the position-specific average signal calculated for each cluster are shown on the right.

**(E)** Bar plots showing the promoter chromatin accessibility status (no ATAC-Seq peak, presence of Common, Gain or Lose peak) for non-expressed genes, non-DEGs and DEGs (all protein-coding) identified in each contrast. Promoters were defined as TSS ± 3 kb. NoPeak: no ATAC-Seq peak was found in any of the gene promoters; Common: at least one Common peak was found; Lose: at least one Lose peak was found; Gain: at least one Gain peak was found; NonExpr: average TPM < 1 in both the conditions of the contrast; NonDEG: the gene is not differentially expressed; DOWN: the gene is downregulated (log2[FC] significantly < 0); DOWN_Strict: the gene is strongly downregulated (log2[FC] significantly < -0.58); UP: the gene is upregulated (log2[FC] significantly > 0); UP_Strict: the gene is strongly upregulated (log2[FC] significantly > 0.58).

**(F)** Visualization of genomic regions encompassing known DE (top panel) and PE (bottom panel) marker gene loci whose promoter hosts a DAR that, respectively, decreases or increases its accessibility during the differentiation to PE, along with tracks relative to (from top to bottom): Bulk RNA-Seq coverage (FPKM values of pooled hESCs, DE, AFE and PE replicates), GENCODE transcripts, ATAC-Seq coverage (RPGC values of pooled DE, AFE and PE replicates), ATAC-Seq peaks (classified based on the AFE vs PE and DE vs PE contrasts) and Vertebrate PhyloP conservation. The image was produced using the UCSC Genome Browser.

**(G)** Visualization of genomic regions encompassing known DE (top panel) and PE (bottom panel) marker gene loci close to non-promoter DARs that, respectively, decrease or increase their accessibility during the differentiation to PE (highlighted by red boxes), along with tracks relative to (from top to bottom): Bulk RNA-Seq coverage (FPKM values of pooled hESCs, DE, AFE and PE replicates), GENCODE transcripts, ATAC-Seq coverage (RPGC values of pooled DE, AFE and PE replicates), ATAC-Seq peaks (classified based on the AFE vs PE and DE vs PE contrasts) and Vertebrate PhyloP conservation. The image was produced using the UCSC Genome Browser.

A

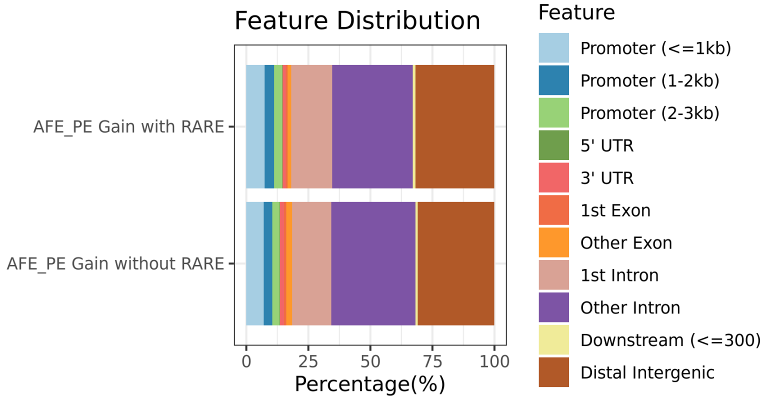
Feature Distribution
AFE_PE Gain with RARE
AFE_PE Gain without RARE
0
25
50
75
100
Percentage(%)
Feature
Promoter (<=1kb)
Promoter (1-2kb)
Promoter (2-3kb)
5' UTR
3' UTR
1st Exon
Other Exon
1st Intron
Other Intron
Downstream (<=300)
Distal Intergenic

B

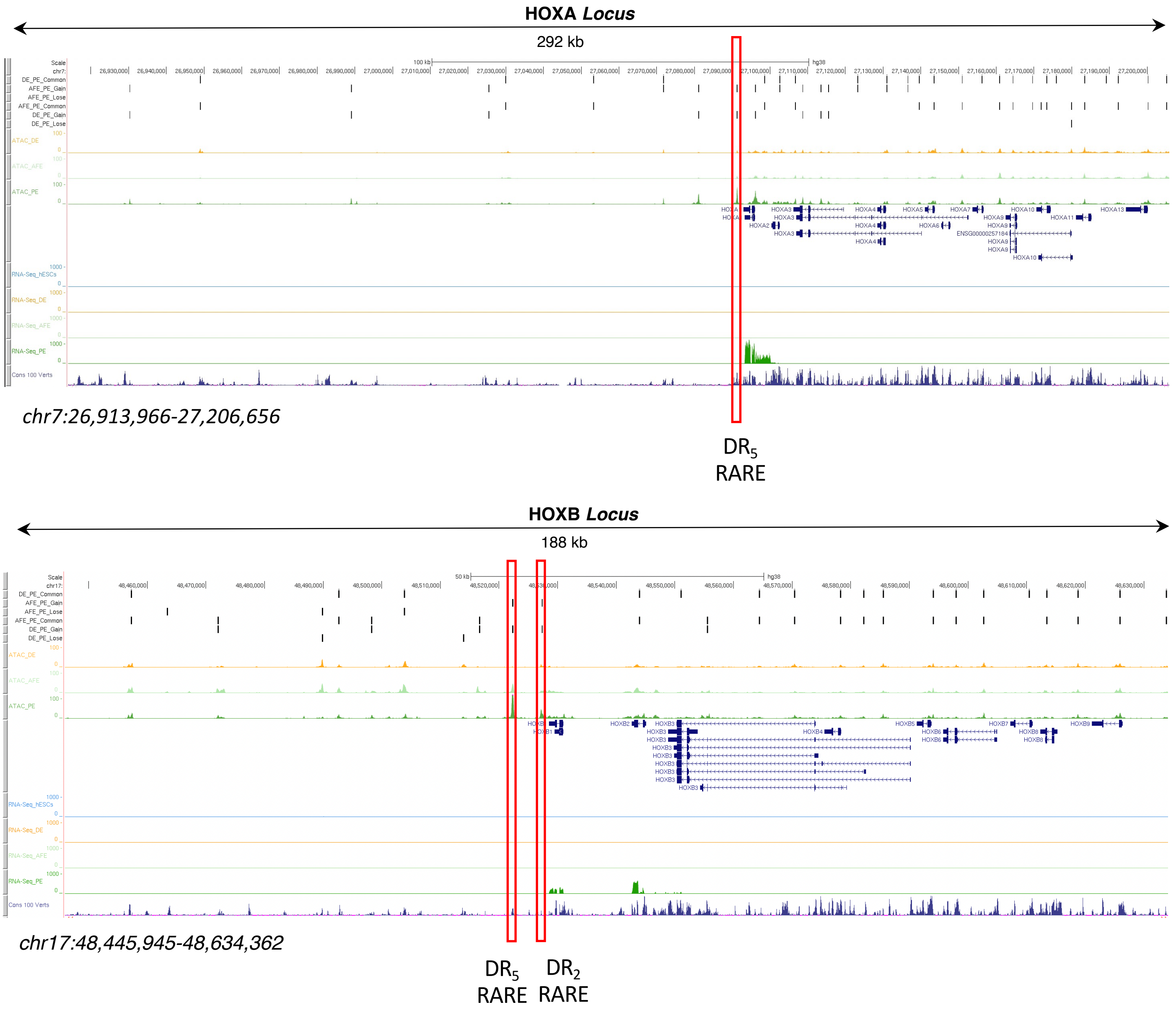
HOXA Locus
292 kb
chr7:26,913,966-27,206,656
DR5
RARE
HOXB Locus
188 kb
chr17:48,445,945-48,634,362
DR5
RARE
DR2
RARE

**Supplementary figure 4:**

**(A)** Bar plot showing the genomic annotation of ATAC-Seq Gain peaks identified in the AFE vs PE contrast that contain or do not contain an enriched RARE FP - i.e. a RARE FP that was found to be enriched among the AFE vs PE Gain peaks (BiFET adjusted p-value < 0.001). Each genomic feature is represented by a specific color shown in the legend

**(B)** Visualization of genomic regions encompassing the HOXA (top panel) and HOXB gene loci, along with tracks relative to (from top to bottom): Bulk RNA-Seq coverage (FPKM values of pooled hESCs, DE, AFE and PE replicates), GENCODE transcripts, ATAC-Seq coverage (RPGC values of pooled DE, AFE and PE replicates), ATAC-Seq peaks (classified based on the AFE vs PE and DE vs PE contrasts), Vertebrate PhyloP conservation and RARE footprints identified in PE. Red boxes indicate the well-known RA-responsive enhancers that control HOXA1 and HOXB1 expression. The image was produced using the UCSC Genome Browser.

### Supplementary figure 5

**A**

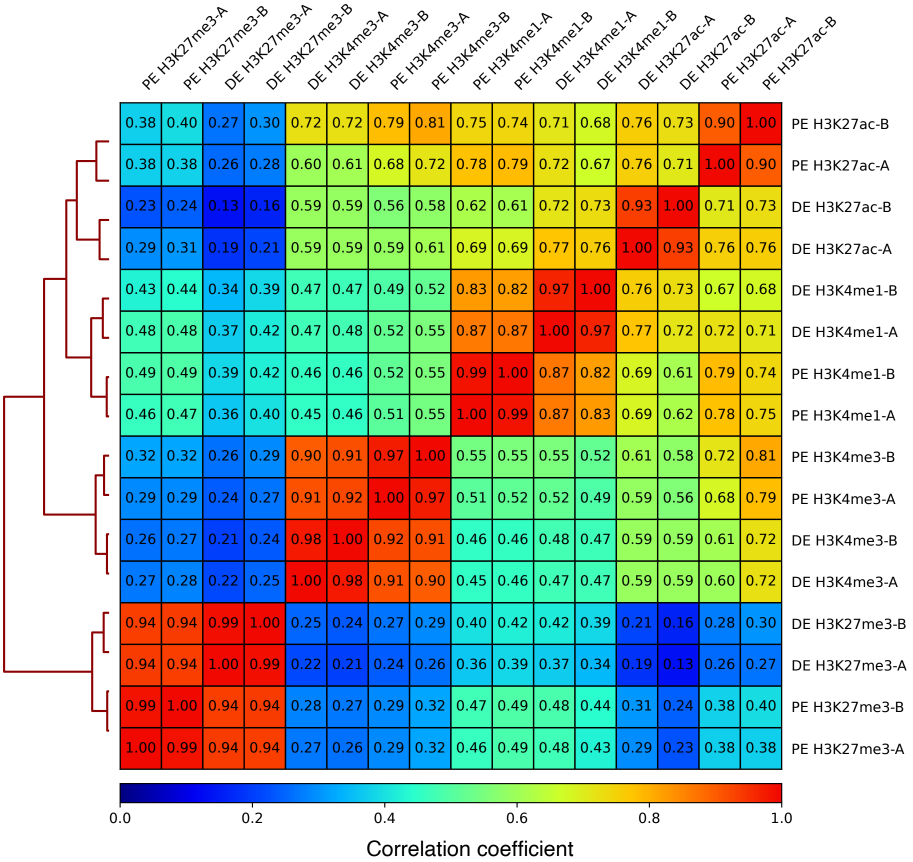


| | PE H3K27me3-A | PE H3K27me3-B | DE H3K27me3-A | DE H3K27me3-B | DE H3K4me3-A | DE H3K4me3-B | PE H3K4me3-A | PE H3K4me3-B | PE H3K4me1-A | PE H3K4me1-B | DE H3K4me1-A | DE H3K4me1-B | DE H3K27ac-A | DE H3K27ac-B | PE H3K27ac-A | PE H3K27ac-B |
|---|---|---|---|---|---|---|---|---|---|---|---|---|---|---|---|---|
| PE H3K27ac-B | 0.38 | 0.40 | 0.27 | 0.30 | 0.72 | 0.72 | 0.79 | 0.81 | 0.75 | 0.74 | 0.71 | 0.68 | 0.76 | 0.73 | 0.90 | 1.00 |
| PE H3K27ac-A | 0.38 | 0.38 | 0.26 | 0.28 | 0.60 | 0.61 | 0.68 | 0.72 | 0.78 | 0.79 | 0.72 | 0.67 | 0.76 | 0.71 | 1.00 | 0.90 |
| DE H3K27ac-B | 0.23 | 0.24 | 0.13 | 0.16 | 0.59 | 0.59 | 0.56 | 0.58 | 0.62 | 0.61 | 0.72 | 0.73 | 0.93 | 1.00 | 0.71 | 0.73 |
| DE H3K27ac-A | 0.29 | 0.31 | 0.19 | 0.21 | 0.59 | 0.59 | 0.59 | 0.61 | 0.69 | 0.69 | 0.77 | 0.76 | 1.00 | 0.93 | 0.76 | 0.76 |
| DE H3K4me1-B | 0.43 | 0.44 | 0.34 | 0.39 | 0.47 | 0.47 | 0.49 | 0.52 | 0.83 | 0.82 | 0.97 | 1.00 | 0.76 | 0.73 | 0.67 | 0.68 |
| DE H3K4me1-A | 0.48 | 0.48 | 0.37 | 0.42 | 0.47 | 0.48 | 0.52 | 0.55 | 0.87 | 0.87 | 1.00 | 0.97 | 0.77 | 0.72 | 0.72 | 0.71 |
| PE H3K4me1-B | 0.49 | 0.49 | 0.39 | 0.42 | 0.46 | 0.46 | 0.52 | 0.55 | 0.99 | 1.00 | 0.87 | 0.82 | 0.69 | 0.61 | 0.79 | 0.74 |
| PE H3K4me1-A | 0.46 | 0.47 | 0.36 | 0.40 | 0.45 | 0.46 | 0.51 | 0.55 | 1.00 | 0.99 | 0.87 | 0.83 | 0.69 | 0.62 | 0.78 | 0.75 |
| PE H3K4me3-B | 0.32 | 0.32 | 0.26 | 0.29 | 0.90 | 0.91 | 0.97 | 1.00 | 0.55 | 0.55 | 0.55 | 0.52 | 0.61 | 0.58 | 0.72 | 0.81 |
| PE H3K4me3-A | 0.29 | 0.29 | 0.24 | 0.27 | 0.91 | 0.92 | 1.00 | 0.97 | 0.51 | 0.52 | 0.52 | 0.49 | 0.59 | 0.56 | 0.68 | 0.79 |
| DE H3K4me3-B | 0.26 | 0.27 | 0.21 | 0.24 | 0.98 | 1.00 | 0.92 | 0.91 | 0.46 | 0.46 | 0.48 | 0.47 | 0.59 | 0.59 | 0.61 | 0.72 |
| DE H3K4me3-A | 0.27 | 0.28 | 0.22 | 0.25 | 1.00 | 0.98 | 0.91 | 0.90 | 0.45 | 0.46 | 0.47 | 0.47 | 0.59 | 0.59 | 0.60 | 0.72 |
| DE H3K27me3-B | 0.94 | 0.94 | 0.99 | 1.00 | 0.25 | 0.24 | 0.27 | 0.29 | 0.40 | 0.42 | 0.42 | 0.39 | 0.21 | 0.16 | 0.28 | 0.30 |
| DE H3K27me3-A | 0.94 | 0.94 | 1.00 | 0.99 | 0.22 | 0.21 | 0.24 | 0.26 | 0.36 | 0.39 | 0.37 | 0.34 | 0.19 | 0.13 | 0.26 | 0.27 |
| PE H3K27me3-B | 0.99 | 1.00 | 0.94 | 0.94 | 0.28 | 0.27 | 0.29 | 0.32 | 0.47 | 0.49 | 0.48 | 0.44 | 0.31 | 0.24 | 0.38 | 0.40 |
| PE H3K27me3-A | 1.00 | 0.99 | 0.94 | 0.94 | 0.27 | 0.26 | 0.29 | 0.32 | 0.46 | 0.49 | 0.48 | 0.43 | 0.29 | 0.23 | 0.38 | 0.38 |

**B**

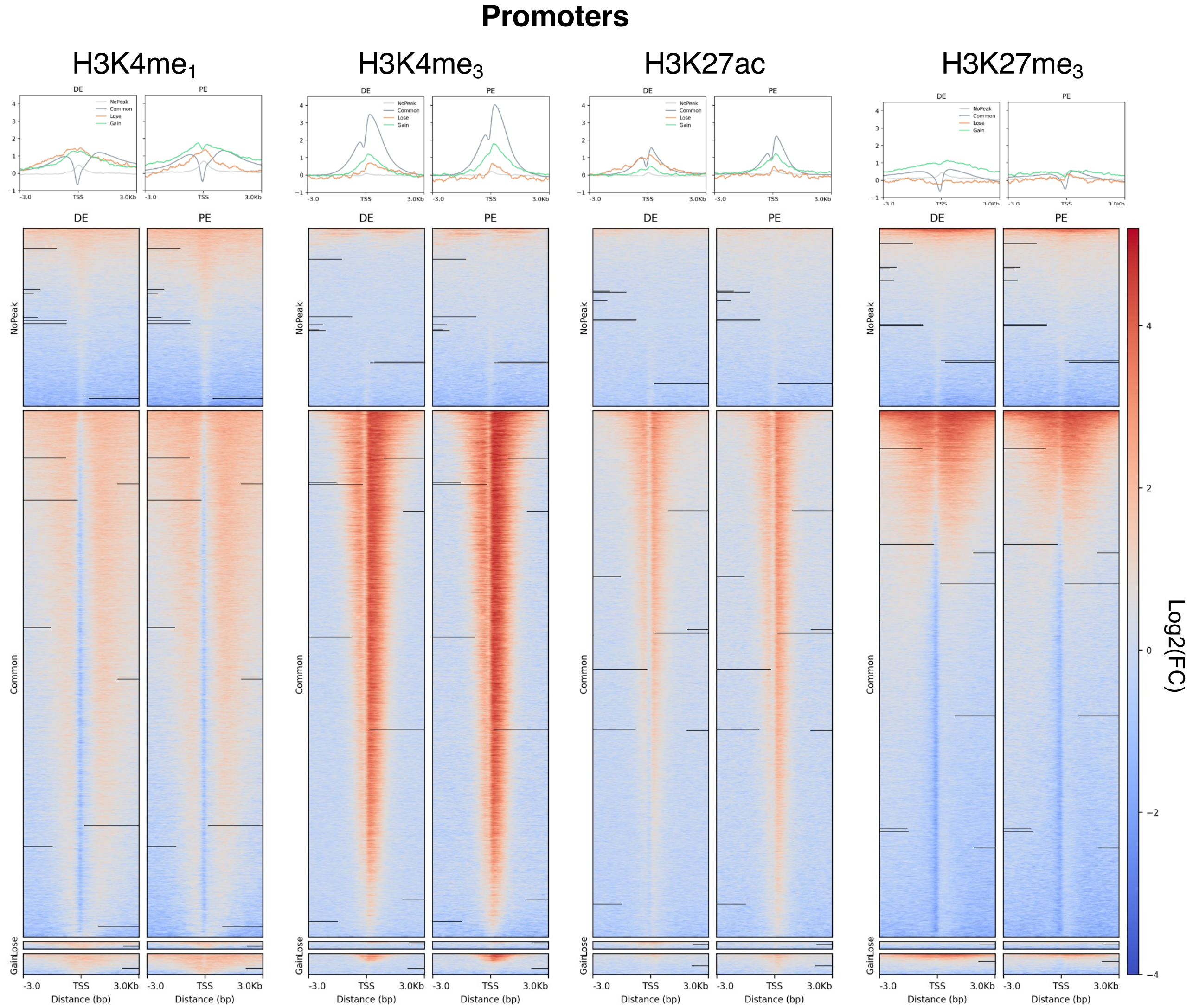

### Supplementary figure 5

C

#### Non-Promoter ATAC-Seq Peaks

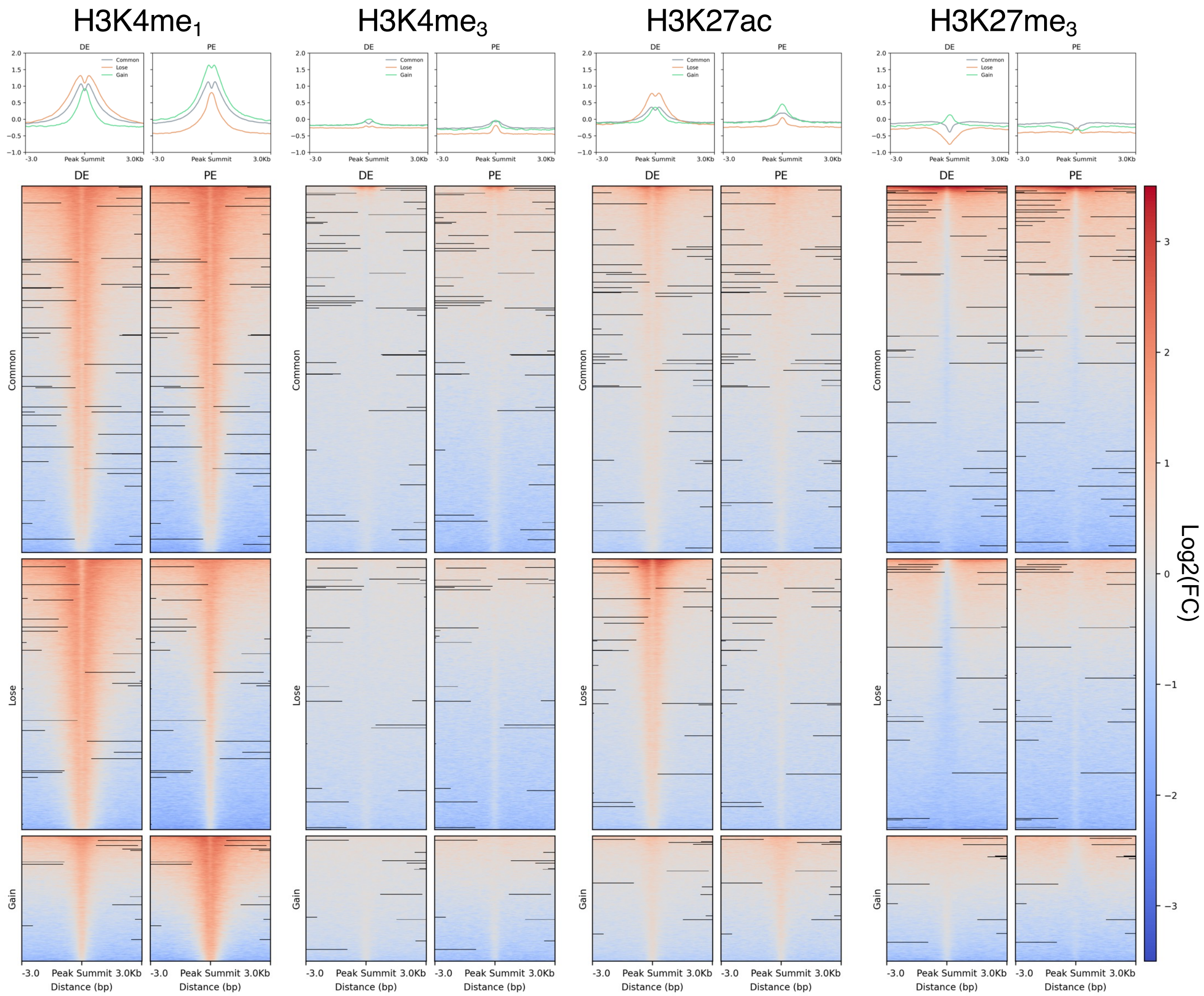


D

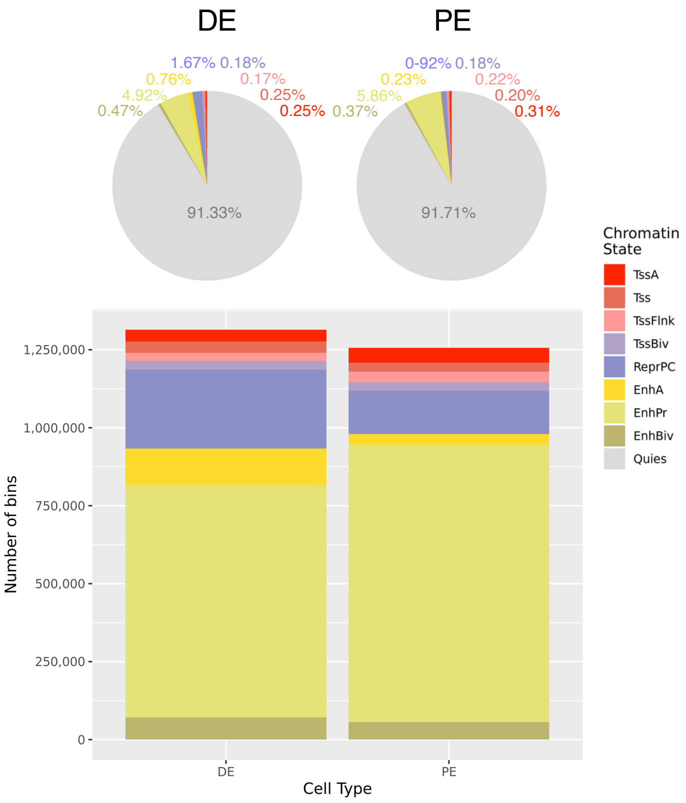


E

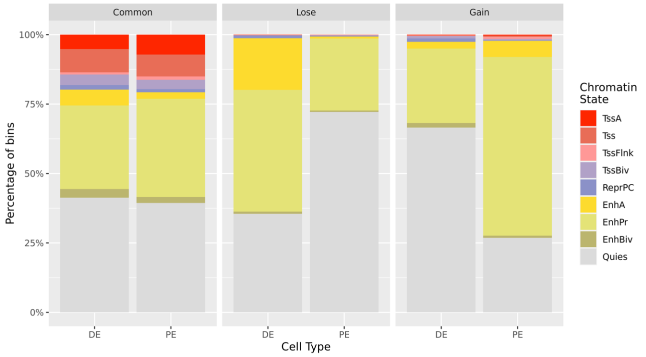


F

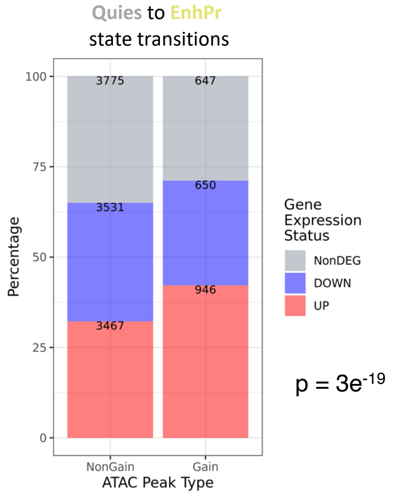

**Supplementary figure 5:**
**(A)** Sample-to-sample Pearson correlation heatmap showing the similarity of DE and PE samples based on their H3K4me3, H3K27me3, H3K4me1, and H3K27ac Histone Mark profiles measured *via* CHIP-Seq of biological duplicates. Sample-to-sample distances were calculated as 1-Pearson correlation of mean Reads Per Genome Coverage (RPGC) scores computed over 10 kb-long genomic bins. Such distances were also used to draw the dendrogram showing the hierarchical clustering of samples.
**(B)** Heatmaps showing DE and PE Histone Mark CHIP-Seq signal in 6 kb-long regions around the TSSs of protein-coding genes, stratified based on the overlap of the TSSs with Common ATAC-Seq peaks or DARs. Signal was calculated on merged replicates as log2-transformed fold change of the RPGC values over the input samples, with a bin size of 50 bp. Summary plots reporting the position-specific average signal calculated for each TSS class are shown above the heatmaps.
**(C)** Heatmaps showing DE and PE Histone Mark CHIP-Seq signal in 6 kb-long regions around the summits of ATAC-Seq peaks localized outside promoter regions (TSS ± 3 kb), stratified based on the differential accessibility between DE and PE. Signal was calculated on merged replicates as log2-transformed fold change of the RPGC values over the input samples, with a bin size of 50 bp. Summary plots reporting the position-specific average signal calculated for each TSS class are shown above the heatmaps.
**(D)** Pie charts (upper panel) and bar plot (bottom panel) showing, respectively, the percentages and the counts of 200 bp-long genomic bins occupied by the different ChromHMM state calls in DE and PE cells. Quiescent states are not represented in the bar plot to better show the differences in the coverage of the other states.
**(E)** Bar plot showing the chromatin state composition in DE and PE cells of 200 bp-long genomic bins that overlap with the Common, Lose and Gain ATAC-Seq peaks identified in the DE vs PE contrast.
**(F)** Bar plot showing, for the DE vs PE contrast, the expression status of the expressed protein-coding genes closest to the genomic regions transitioning from a Quies state in DE to an EnhPR state in PE, stratified based on the overlap with Gain ATAC-Seq peaks. Nearest genes were searched within a 50 kb range. The reported p-value was calculated by comparing the proportions of upregulated genes between the two classes of genes *via* Fisher's exact test. NonDEG: the gene is not differentially expressed; DOWN: the gene is downregulated (log2[FC] significantly < 0); UP: the gene is upregulated (log2[FC] significantly > 0); NonGain: the genomic region does not overlap with a Gain peak summit; Gain: the genomic region overlaps with a Gain peak summit.

Supplementary figure 6

A

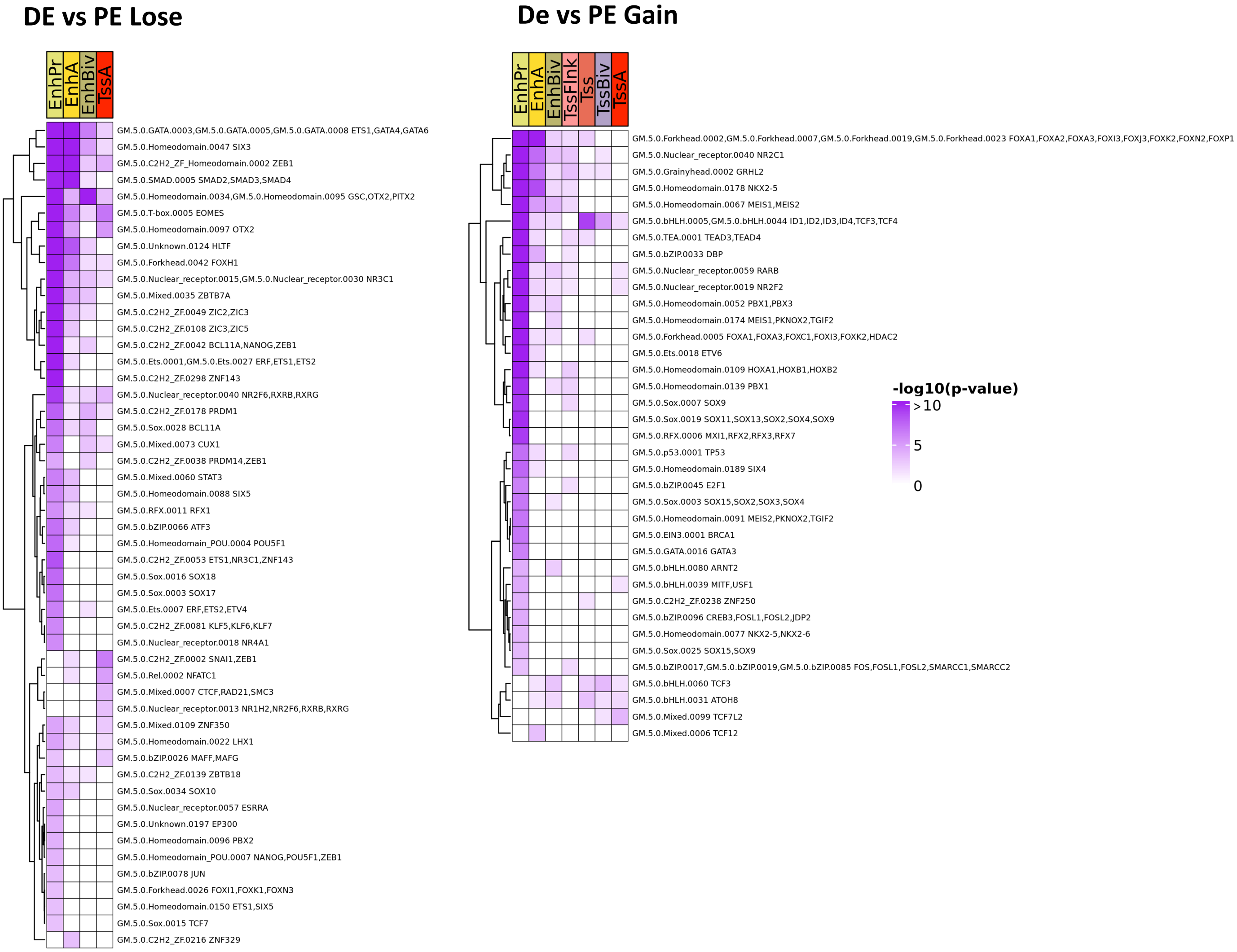


B

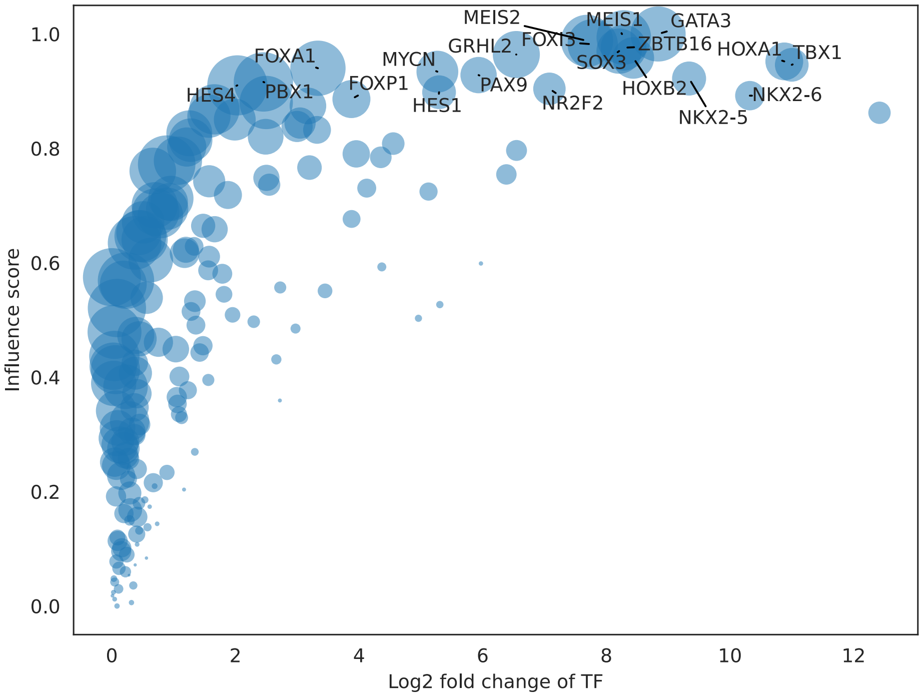

**Supplementary figure 6:**
**(A)** Heatmaps showing TF FPs enriched in DE vs PE Lose (left panel) and Gain (right panel) DARs, stratified based on their DE and PE chromatin state, respectively. For the analysis of Lose DARs, only transitions enriched in Lose peaks and downregulated TFs with an average DE TPM > 5 were used; for the analysis of Gain DARs, only transitions enriched in Gain peaks and upregulated TFs with an average PE TPM > 5 were employed. Each row refers to one or more GimmeMotifs vertebrate motifs having identical enrichment profile and the corresponding TFs. Redundant motifs were excluded for better visualization. Cell color intensities are proportional to the –log10 transformed BiFET p-value; only motifs having p-value < 0.001 in at least one DAR class are reported. **(B)** Bubble plot showing the Ananse TF sumScaled influence scores for the transition from DE to PE, along with the log2(FC) calculated for the DE vs PE contrast. Bubble size is proportional to the number of direct targets. The top 20 TFs based on influence score are labelled with their gene name.
